# PHENOTYPIC CHARACTERIZATION AND BACTERIAL INTERACTIONS OF THE COMPLEX MICROBIAL POPULATION IN URINARY DOUBLE-J CATHETERS

**DOI:** 10.1101/2024.12.04.626871

**Authors:** Juan Vicente Farizano, Emilia Castagnaro, Julián Arroyo-Egea, Juan Daniel Aparicio, Cecilia Vallejo, Elvira María Hebert, Lucila Saavedra, Viviana Andrea Rapisarda, Josefina María Villegas, Mariana Grillo-Puertas

**Affiliations:** Instituto Superior de Investigaciones Biológicas (INSIBIO), CONICET-UNT, and Instituto de Química Biológica “Dr. Bernabé Bloj”, Facultad de Bioquímica, Química y Farmacia, UNT. Chacabuco 461, T4000ILI-San Miguel de Tucumán, Argentina; Planta Piloto de Procesos Industriales Microbiológicos (PROIMI-CONICET). Tucumán, Argentina; Hospital Eva Perón, Ministerio de Salud Pública de Tucumán; Centro de Referencia para Lactobacilos (CERELA-CONICET), Tucumán

**Keywords:** Uropathogens, catheters, biofilm, virulence, bacterial interaction, antibiotic resistance

## Abstract

Catheter-associated urinary tract infections (CAUTIs) are a major cause of morbidity in healthcare settings, often linked to the colonization of catheters by diverse bacterial species. This study aimed to characterize 27 clinical uropathogenic isolates obtained from double-J catheters through phenotypic and genotypic assays. The isolated species included, *Staphylococcus* spp., *Enterococcus faecalis*, *Klebsiella pneumoniae*, *Escherichia coli* and *Bacillus* spp. These isolates were evaluated for biofilm formation, production of extracellular components such as cellulose and amyloid-like fibers, motility, hemolytic capacity, siderophore production, stress tolerance, and antibiotic resistance profiles. The findings revealed that *E. coli*, *S. aureus*, *B. subtilis* isolates exhibited strong biofilm formation and robust extracellular matrix production, features that contribute to their persistence in catheter environments. Acid pH and oxidative stress tolerance results indicated a differential strain response even within the same genus. Additionally, multidrug resistance (MDR) was observed in several isolates, especially in *K. pneumonia* and *E. faecalis*, correlating with their biofilm-forming capacities. Co-cultures studies demonstrated synergistic interactions between the co-isolated pair, particularly with *Bacillus* spp. enhancing biofilm formation in static condition. The interaction results indicate the importance of investigating the potential clinical relevance of *Bacillus* spp., even though they are not traditionally considered human pathogens. These results provide critical insights into the pathogenic mechanisms and survival strategies of several uropathogens in CAUTI, emphasizing the need for targeted therapeutic interventions to prevent recurrent infections and manage antibiotic resistance.

**Importance:** Catheter-associated urinary tract infections (CAUTIs) are a persistent challenge in healthcare due to their association with biofilm-forming bacteria that resist treatments. To investigate the complexity of microbial population of CAUTI, we provide an in-depth analysis of the diverse range of bacterial strains isolates from the devices. This study highlights the role of both well-known pathogens and less traditionally recognized species, such as *Bacillus* spp., in these infections. By revealing that bacterial interactions analyzed here enhance biofilm formation, this research underscores the need to reconsider the clinical importance of *Bacillus* spp. in polymicrobial settings. This knowledge is critical for designing more effective therapeutic interventions to mitigate biofilm-related complications in medical devices.

## Introduction

Urinary tract infections (UTI) affect a large number of people, with their prevalence varying according to age group, sex, and the presence of risk factors (1–4). The high incidence of UTI, together with the increased recurrence rate and the complicated forms that may arise significantly impact on the cost of medical treatment, resulting in billions of dollars in health care costs (5). The primary cause of UTI is the invasion of the urinary tract by microorganisms known as uropathogens (UP). The main etiologic agent for UTI cases in humans is uropathogenic *E. coli* (UPEC), responsible for up to 80% of all infections (6), while microorganisms such as *Staphylococcus* spp., *Proteus mirabilis*, *Proteus. vulgaris*, *Klebsiella* spp., *Enterococcus faecalis* and *Pseudomonas aeruginosa* account for the remaining 20% (7, 8).

The risk of developing a UTI significantly increases with the use of indwelling devices, since it disrupts host defense mechanisms and facilitates access of UP to the bladder or to the upper urinary tract (9). The double-J stent is a commonly used device in urological surgery, and presents dual functions by providing support and drainage (10–12). Catheter-associated urinary tract infections (CAUTI) are a significant concern in healthcare settings, as can cause increased morbidity, long hospital stays, and high healthcare costs (13). CAUTI-causing microorganisms originate from the patient own microbiota, often modified by antibiotic pressure, and cross-transmission by healthcare workers. The surface of urinary catheters offers a favorable environment for the colonization and growth of a large number of microorganisms (14–16). Therefore, exploring the clinical characteristics of CAUTI, and the features of the involved pathogenic bacteria is central to treat these infections. An additional problem with CAUTI is their diagnostic difficulties, as colonizing bacteria may not appear in urine or prostatic secretion samples (17, 18).

Bacterial populations have the capacity to adapt quickly to changes in their environmental surroundings (19, 20). The ability of UP strains to cause different types and severity of diseases depends on the expression of multiple virulence traits, such as biofilm formation, motility, hemolytic capacity, expression of several virulence genes related to adhesin production, toxins, siderophores, and secretion systems, among others (21, 22). UP are often found in sessile form of life on the intra- and/or extraluminal surfaces of urinary catheters removed from patients with UTI (23). Over the years, several reports have shown that most biofilms formed on long term-catheters are polymicrobial, with an average of 2-5 microbial species isolated per catheter, predominantly consisting of Gram-negative bacteria (24–28). UP isolates from polymicrobial biofilms in CAUTI patients exhibited significant resistance to commonly used antibiotic agents, thereby increasing the risk of developing severe infections (29, 30). In the context of the urinary tract, the presence of multiple microorganisms in a midstream urine sample are often suspected to be the result of periurethral or vaginal microbiota contamination (31, 32). In fact, polymicrobial samples may also be labeled as negative culture if they lack a single dominant species, or they may be categorized as having secondary pathogens or doubtful pathogens depending on the level of colonization. Therefore, a more comprehensive understanding of UP virulence and the often-underestimated complexity of polymicrobial infections in the context of CAUTI is essential for improving patient care and minimizing associated morbidities. The aim of this work was to investigate the virulence-associated phenotypic traits and antimicrobial resistance of bacteria isolated from double-J catheters, as well as to explore the interactions within mixed biofilms formed by co-isolated bacterial pairs. Here, an exhaustive study of the multiple virulence factors expressed by CAUTI isolated bacteria and their interaction were carried out. Such knowledge would be important for managing catheter-associated infections that impact in patient outcomes and healthcare systems.

## MATERIAL AND METHODS

### Catheters processing and UP isolation

Samples were taken from 8 double-J type urinary catheters provided by a specialized private urological center in the city of San Miguel de Tucumán, Argentine. In all cases, 30-60 days urinary catheters were collected from adult male and female patients with negative urine culture, with or without a chronic pathology. Each catheter was processed under sterile conditions to remove adherent and non-adherent bacteria, as reported by Mandakhalikar et al. (2018). Briefly, to extract unattached cells, 4 washes with physiological saline solution (PS) were performed. To extract adhered bacteria from catheters, the vortexing–sonication–vortexing (V-S-V) method was used (33). Both unattached and attached bacteria were concentrated by centrifugation for 10 min at 8000 rpm. Then, cells were resuspended in PS and seeded with Digralsky spatula on CLED-agar plates. The plates were incubated during 24-48 h at 37°C to evaluate microbial growth. Colonies with different morphologies were purified by the stretch marks method in CLED medium. Pure isolates of potential UP were preserved for subsequent identification.

### Identification of UP isolates

In order to identify the obtained UP isolates, genomic DNA extraction was performed using the organic solvent extraction method reported by Sambrook et al (1989) (34). Quantification of the extracted genomic DNA was carried out using a Nano-Drop spectrophotometer (UV/Vis nano spectrophotometer, Nabi). Then, 16S rRNA gene amplification was carried out and sequenced to identify the genus of the obtained UP isolates. Also, identification of UP at species level was carried out according to the standard methods and recommendations of the International Committee of Clinical Laboratories (35). Standard biochemical tests and confirmation by MALDI-TOF mass spectrometry techniques were also used for identification (Bruker Daltonics) (36, 37).

### Culture conditions

Isolates were routinely grown under aerobic conditions, in BHI medium at 37°C with shaking (180 rpm), or in static growth at 37°C on CLED-agar plates. Bacterial growth was monitored by measuring absorbance at 600 nm (A_600nm_) to determine turbidity in liquid medium, or by the appearance of isolated colonies in solid medium. MacConkey, LB, M63, BHI and urine media were used, depending on the assay. Human urine media was prepared according to Eberly et al. (2020) (38). Briefly, urine was collected in equal volumes from male and female healthy volunteers and filtered through a 0.22 µm filter prior to use. Healthy volunteers were classified as individuals who are urologically asymptomatic and not menstruating, and who have not taken antibiotics in the last 90 days.

### Biofilm formation and quantification

Biofilm formation was assayed according to the ability of cells to adhere and grow in 96-well polystyrene plates in different culture media (20). Briefly, 24 h cultures of all isolates were washed in PS and subsequently diluted to A_600nm_ = 0.1 in the corresponded media. Suspensions were loaded in multiwell plates and incubated under static conditions at 30°C for the indicated times in each assay. Then, unattached (planktonic) cells were removed, and the wells were washed three times with distilled water. Quantification of adherent cells or biofilm biomass was performed as follows: 200 μL of a 0.1% crystal violet solution was added to each well and the plates were incubated at room temperature for 15 min in the dark; then, the wells were washed three times with distilled water to remove crystal violet excess. The dye staining the adhered cells (biofilm) was solubilized with 200 μl of 95% ethanol and A_595nm_ was measured on a microplate reader (SpectraMax Plus384, USA.). Each condition was performed in quadruplicate and, each experiment was performed no less than five times.

UP isolates were classified according to their biofilm formation capacity after 72 h, following the Stepanovic criteria (39). The A_595nm_ cut-off value (**A_595nm_C**) was defined as the average value of the blank absorbance plus three standard deviations. Blank was the non-inoculated medium. Hence, the following classification criteria were established: A_595nm_ ≤ **A_595nm_C**, non-biofilm producers (-); **A_595nm_C** < A_595nm_ ≤ 2 fold **A_595nm_C**, weak biofilm producers (+); 2 fold **A_595nm_C** < A_595nm_ ≤ 4 fold **A_595nm_C**, moderate biofilm producers (++); A_595nm_ > 4 fold **A_595nm_C**, strong biofilm producers (+++), and A_595nm_ > 8 fold **A_595nm_C**, robust biofilm producers (++++).

### Colony morphotypes: amyloid-like fibers production and cellulose production

Colony morphology and the ability to bind certain dyes may serve as an indicator of several physiological and metabolic microbial states (40). To analyze colony morphology, culture plates supplemented with Congo Red (CR) and Brillant Blue (BB) were used as described by Da Re and Ghigo (2006) (41). Briefly, BHI cultures of all isolates were grown overnight (ON) at 37°C, washed and diluted to OD_600nm_ = 0.1 in PS. Subsequently, 5 µl of these suspensions were spotted on LB-agar plates (low salt) supplemented with CR 40 µg/ml and BB 20 µg/ml. Plates were incubated at 30°C for 96 h and monitored every 24 h. Colony phenotypes analysis was performed for every bacterial genus, according to the previously reports. For *E. coli* and *K. pneumoniae*, the analysis was according to Bokranz et al. (2005): *ras* (red and smooth colonies) and *pas* (pink and smooth colonies) morphotypes (42). For *Staphylococcus* spp. isolates, the results were interpreted as reported by Arciola et al. (2002): reddish-black colonies with a rough and dry consistency were considered biofilm-related extracellular matrix-producing strains (43). Morphology of *E. faecalis* strains was analyzed as reported by Torres-Rodriguez et al. (2020), where strains with black and rough colonies are considered as biofilm producers and strains with red or white colonies as non-producers (44). For *Bacillus* spp., bacteria that bind Congo Red dye, have previously been classified as functional bacterial amyloid fibers (45). The presence of amyloid-like fiber was denoted as +.

Cellulose production was assayed as described by White et al. (2006) (46). For this, BHI cultures of each isolate were grown ON at 37°C, washed and diluted to OD_600nm_ = 0.1 in PS. Subsequently, 5 µl of the isolates were spotted on LB-agar plates supplemented with Calcofluor White dye 50 μg/mL (Sigma-Aldrich). Plates were incubated at 30°C for 96 h and monitored every 24 h. Cellulose production was qualitatively assessed by observing colonies fluorescence when irradiated with UV light. Fluorescent colonies were denoted as +.

### Motility assay

Bacterial motility was assessed according to Ulett et al (2006), with minor modifications (47). Briefly, ON cultures on BHI were washed and diluted to OD_600nm_ =0.1 in PS. Suspensions were then seeded onto semi solid LB-agar plates (0.3%) by using a sterile toothpick. Plates were incubated at 30°C and the colony diameter was evaluated over time. Isolates with colony diameter between 0.5 and 1 cm were denoted as + and more than 1 cm, ++.

### Analysis of hemolytic capacity

The isolates hemolytic activity was assessed on blood agar plates according to Gerhardt et al (1994) (48). Briefly, isolates were grown ON in BHI medium, then the cells were washed and diluted to OD_600nm_ = 0.1 in PS. Aliquots of the bacterial suspensions were seeded on plates of blood agar medium and incubated for 24 h at 30°C. A β-hemolytic reaction involves complete lysis of red blood cells, causing a clear area on the agar surrounding the colony, called total hemolysis. On the other hand, an α-hemolytic reaction occurs when the hemoglobin in red blood cells is reduced to methemoglobin, causing a greenish tint in the agar surrounding the colonies. Finally, the absence of hemolysis or discoloration is termed γ-hemolysis (49).

### Siderophores production assay

The siderophores production was determined by a qualitative technique, adapted in our laboratory, based on the color change of Cromoazurol S. Strains were cultured ON in M9 minimal medium supplemented with 0.2% glucose with shaking at 37°C. 5 µL aliquots of each cell culture was seeded on M9 minimal medium supplemented with 0.2% glucose plates and incubated at 37°C for 48 h. Then, an overlaid of semisolid CAS medium (Cromoazurol 60.5 mg, piperazine acid 72.9 mg, FeCl_3_ 1 mM, dissolved in HCl at 10 mM, 10 mL of the solution, for 1 liter) was poured onto the grown plate. Those isolates that are capable of producing siderophores presented a color change of the CAS medium from blue to yellowish around the colony.

### Hydrogen peroxide, acid and human serum tolerance assays

Stressors susceptibility was carried out according to Shea et al. (2022) (50). Briefly, bacterial cultures were incubated ON in BHI medium at 37°C. Cultures were normalized to 0.5 McFarland (∼ 1.5 x 10^8^ CFU/mL) (McFarland J, 1907) in 1 mL of either BHI, BHI containing fresh 0.2% H_2_O_2_ or BHI buffered to pH 7, 5, or 2.5. Samples were immediately vortexed and incubated during 15 or 60 min at room temperature for H_2_O_2_ assay, and 1 h at 37°C with aeration for acid tolerance assay. At each time point, suspensions were serially diluted in PS to CFU/mL determination. For human serum tolerance, ON BHI cultures (1 mL) were pelleted by centrifugation and 0.5 McFarland suspensions were prepared in sterile PS. Suspensions were diluted 1:10 in sterile PS. Ten microliters of the diluted solution was added to 190 mL of either 100% human serum or 100% heat-inactivated human serum (Innovative Research catalog no. ISER10ML). The mix was incubated for 1 h at 37°C, and then the number of CFU/mL was calculated by serially diluting the bacteria-serum suspension and plated on CLED-agar medium. CFU/mL of the bacterial inoculum was calculated by serially diluting the bacterial suspension and plated on CLED-agar medium.

### Antibiotic susceptibility

Antibiograms were performed for each UP using the disc diffusion method on Müller-Hinton agar (MH-agar) (BB-NCIPD Ltd.) (35). The antimicrobials choice is based on national and international recommendations for urinary infection in adult, inpatient and outpatient. The ATBs used for Gram-negative and Gram-positive bacteria in this work were: Norfloxacin (NOR) 10 μg, Gentamicin (GEN) 10 μg, Nitrofurantoín (NIT) 300 μg, Ampicilin Sulbactame (AMS) 10/10 μg, Trimethoprim/Sulfamethoxazole (SXT) 1.25/ 23.75 μg, nalidixic acid (NA) 30 μg, Ciprofloxacin (CIP) 5 μg, Ampicillin (AMN), Amikacin (AKN) 30 μg, Ceftazidime (CAZ) 30 μg, Cefuroxime (CXM) 30 μg and Imipenem (IMP) 10 μg. In addition, ATB specifically form Gram-positive *cocci* were also tested: Oxacillin (OXA) 1 μg, Erythromicin (ERY) 15 μg, Clindamycin (CLI) 2 μg, Vancomycin (VAN) 30 μg and cefoxitin (FOX) 30 µg. Briefly, each isolate was replicated on individual CLED-agar plates. Colonies were resuspended in 5 mL of PS, and suspensions were adjusted to 0.5 on the McFarland scale. Standardized suspensions were seeded with a sterile swab on MH-agar plates in order to obtain a cell lawn. ATB discs were placed on the lawn carefully with sterile forceps, and plates were incubated ON at 37°C. Inhibition halos were measured and interpreted according to the following references (35): Enterobacteriaceae: ≤ 13 mm, Resistant; 14-16 mm, Intermediate; ≥ 17 mm, Susceptible. *Staphylococcus* spp.: ≤ 12 mm, Resistant; 13-14 mm, Intermediate; ≥ 15 mm, Susceptible. *Enterococcus* spp.: ≤ 14 mm, Resistant; ≥ 15 mm, Susceptible. *Bacillus* spp.: ≤ 10 mm, Resistant; 11-12 mm, Intermediate; ≥ 13 mm, Susceptible. MDR was defined as acquired non-susceptibility to at least one agent in three or more antimicrobial categories, XDR was defined as non-susceptibility to at least one agent in all but two or fewer antimicrobial categories (i.e. bacterial isolates remain susceptible to only one or two categories) and PDR was defined as non-susceptibility to all agents in all antimicrobial categories (51)

### Detection of Genes Encoded Virulence Factors

The screening of the presence/absence of virulence-associated genes was carried out by PCR. For each bacterial genus, the following genes were tested: for *E. coli,* haemolysin (*hlyA*), cytotoxic necrotising factor (*cnf1*), fimbria type I regulator (*fimB*), type 1 fimbria (*fimA*), fimbria P (*papA*) and iron uptake related genes (*iroN* and *iutA*); for *K. pneumoniae,* haemolysin (*hlyA*), type 1 fimbrial subunit (*fimA*), the type 3 fimbrial adhesin (*mrkD*)*, enterobactin* (*entB*)*, and* an outer membrane porin (*ompK36*); for *E. faecalis,* gelatinase (*gelE*), sortase-type enzyme (*srt*), bacterial adherence protein (*efaA*), collagen adhesin protein (*ace*), cytolicin production activator (*cylA*); for *Staphylococcus* spp, staphylococcal enterotoxin A (*sea*), slime production related protein (*ica*), arginine metabolism related protein (*argB*, *argC*), lipase (*gehC*), fibrinogen binding protein (*sdrG*) and for *Bacillus* spp., bacillus enterotoxins (*entB, entA*), pore-forming toxin (*cytK*), haemolysin (*hlyIII*), sporulation related -ATPase subunit (*clpC*), capsule synthesis protein (*capA*). PCR reactions were performed using the primers and parameters described in **Table S1**. The PCR reaction mixture (25 μL total) consisted of 12.5 μL of 2× SYBR Green PCR Master Mix (Bio-Rad), 300 nmol each of forward and reverse primers, and 10 ng of genomic DNA as the template. The amplification protocol included an initial denaturation at 95°C for 4 minutes, followed by 35 cycles of denaturation at 95°C for 15 seconds and annealing/extension at 56°C for 30 seconds. Fluorescence signals were recorded at the end of each extension step, and a melt curve analysis was performed to verify product specificity. To confirm amplification, 5 μL of the PCR product was analyzed on a 2.0% agarose gel alongside a 100 bp DNA ladder.

### Bacterial interaction assays: colony interaction assay, liquid medium interaction and mixed biofilm formation

Bacterial interaction was evaluated in co-isolated pairs obtained simultaneously from the same catheter. Four co-isolated pairs were selected, consisting of *Bacillus* strains with other UP: *E. faecalis* Ef5/*B. pumilus* Bp1, *S. epidermidis* Se3/*B. subtilis* Bs1, *S. epidermidis* Se4/*B. subtilis* Bs2 and *K. pneumoniae* Kp2/*B. megaterium* Bm2. For colony interaction assays, a working protocol developed in the laboratory was followed. Briefly, bacteria were grown overnight (ON) at 37°C in BHI liquid medium with shaking (150 rpm). Then, 5 µL aliquots of the cultures were taken and placed on the surface of a solid BHI-agar medium plate, following the scheme of one central bacterium and four concentric bacteria at different distances. Bacteria (central or peripheral) were seeded simultaneously or in a deferred manner (24 h after the growth of the central colony). The interaction was analyzed over a period of 5 days at 30°C. For liquid media co-culture assays, a previously described protocol was carried out (52–54). Briefly, isolates were grown ON at 37°C in BHI, harvested by centrifugation and adequately diluted in BHI medium to an OD_600nm_=0.1. Monospecies or mixed cultures were grown under shaking conditions at 37°C for different times, and viability measured, using appropriate media and ATB to achieve single species identification in the interaction. For mono or multispecies biofilm assays, isolates were grown at 37°C in BHI and adequately diluted to a OD_600nm_=0.1. Cells of different bacteria (alone or in co-culture) were grown in multiwell plates during 24 h at 30°C. After incubation under static conditions, the non-adherent fraction (planktonic cells) was removed and total biofilm biomass was quantified using the crystal violet technique (20). Non adhered fraction containing mixed culture was quantified by UFC/mL using appropriate media as mentioned before. To quantified microorganisms in the biofilm biomass, adhered cells were washed three times with sterile PS and incubated with Triton 0.1% during 10 min. Biofilm cells were extracted by scraping vigorously with a sterile tip. These bacteria were resuspended in sterile PS and quantified by determining CFU/mL using appropriate media as mentioned before (53).

### Statistical analysis

The statistical analyses were conducted using R (version 4.4.1) and RStudio (version 2024.09.0+375). For all inferential tests, a *p*-value of less than 0.05 was considered statistically significant. Normality and homoscedasticity were assumed for the data. IN cases where necessary, a log-transformation (log1p) was applied to stabilize variances and improved normality, ensuring that the assumptions of parametrics tests were met. A two-way ANOVA was performed, followed by post-hoc comparisons using the Fisher method where appropriate. Additionally, a Principal Components Analysis (PCA) was carried out to identify clusters and explore relationships among the variables. A biplot was generated to visualize both the variables and clusters, providing a clear graphical representation of the data’s structure and groupings.

## RESULTS AND DISCUSSION

### Isolation and identification of microorganisms

A total of 27 microorganisms were isolated from 8 catheters **(Table 1)**. Monobacterial isolates were obtained from 3 catheters (catheters 1, 4, and 5), whereas multiple bacterial species were recovered from the remaining catheters, confirming the polymicrobial etiology of CAUTI (18, 55, 56). The predominant bacterial species were *Bacillus* spp., including *B. pumilus*, *B. subtilis*, and *B. megaterium* (7 isolates, 27%), followed by *Staphylococcus* spp., including *S. aureus* and *S. epidermidis* (1 and 6 isolates, respectively, 27%). *E. faecalis* (5 isolates, 19%), *K. pneumoniae* (4 isolates, 15%), and *E. coli* (3 isolates, 12%) (**Table 1**). It is worth to mention that all patients had the catheters in place for a time period of 1 to 3 months, and prior to extraction, all presented negative urine cultures (UC). Additionally, prior catheter extraction, antibiotic therapy was administered to patients. Even though, adhered and non-adhered bacteria were isolated from the devices. The identified species (except for *Bacillus* spp.) are in agreement with prior literature concerning bacterial species in CAUTI, described as a common uropathogen in UTI (57–60). Although only 12% were UPEC isolates, this bacterium is the most common pathogen in both community-acquired and nosocomial urinary tract infections, being responsible for approximately 24-39% of CAUTIs (61). It is widely known that *K. pneumoniae* is an opportunistic pathogen commonly distributed in the perineum, which can easily colonize the urinary tract and medical devices (60). *E. faecalis* establishes a symbiotic relationship with other UP, end is frequently found in CAUTI contexts. (62). In addition, *S. aureus* and *S. epidermidis*, normally present on the skin surface, can enter the urinary tract during catheter placement (18, 63, 64). Often, the cause of this colonization may be due to the fact that catheterization and its manipulation promote ascending infections due to the presence of commensal microorganisms in the urethra, surrounding skin, or potential contamination resulting from healthcare personnel manipulation (55, 65). Notably, it was observed that several bacterial isolates adhered to different catheters belong to the *Bacillus* genus. It is well known that human infections caused by *Bacillus* spp., except for anthrax, are rarely reported in the literature. The main reason for this inadequate reporting by clinical laboratories could be explained by the fact that most of these bacteria are saprophytic, and their isolation from human samples is often ignored and considered as laboratory contaminants (66, 67). However, some reports described that *Bacillus* spp. have been isolated from bacteremia, endocarditis, wounds, respiratory, urinary and gastrointestinal tract infections, food poisoning, and meningitis (66, 67). Therefore, the identification of aerobic gram positive spore-forming bacilli at species level, the evaluation of their pathogenic potential and the interpretation of susceptibility ATB tests of these bacilli, would be relevant to elucidate their potential role in the mentioned infection (18, 68).

**Table 1.** Clinical isolates obtained from double-J catheters.

| **Name** | **Identification** | **Source** |
| --- | --- | --- |
| **Ec1** | *Escherichia coli* | Catheter 2 - Adhered |
| **Ec2** | *Escherichia coli* | Catheter 2 – Non Adhered |
| **Ec3** | *Escherichia coli* | Catheter 2 – Non Adhered |
| **Kp1** | *Klebsiella pneumoniae* | Catheter 8 – Adhered |
| **Kp2** | *Klebsiella pneumoniae* | Catheter 8 – Adhered |
| **Kp3** | *Klebsiella pneumoniae* | Catheter 8 – Adhered |
| **Kp4** | *Klebsiella pneumoniae* | Catheter 8 – Adhered |
| **Ef1** | *Enterococcus faecalis* | Catheter 2 - Adhered |
| **Ef2** | *Enterococcus faecalis* | Catheter 2 - Adhered |
| **Ef3** | *Enterococcus faecalis* | Catheter 2- Adhered |
| **Ef4** | *Enterococcus faecalis* | Catheter 2 – Non Adhered |
| **Ef5** | *Enterococcus faecalis* | Catheter 3 - Adhered |
| **Ef6** | *Enterococcus faecalis* | Catheter 3 - Adhered |
| **Sa1** | *Staphylococcus aureus* | Catheter 1 - Adhered |
| **Se1** | *Staphylococcus epidermidis* | Catheter 4 - Adhered |
| **Se2** | *Staphylococcus epidermidis* | Catheter 5 - Adhered |
| **Se3** | *Staphylococcus epidermidis* | Catheter 6 - Adhered |
| **Se4** | *Staphylococcus epidermidis* | Catheter 6 – Non Adhered |
| **Se5** | *Staphylococcus epidermidis* | Catheter 7 - Adhered |
| **Se6** | *Staphylococcus epidermidis* | Catheter 7 –Non Adhered |
| **Bm1** | *Bacillus megaterium* | Catheter 4 – Non Adhered |
| **Bm2** | *Bacillus megaterium* | Catheter 8 – Adhered |
| **Bp1** | *Bacillus pumilus* | Catheter 3 - Adhered |
| **Bs1** | *Bacillus subtilis* | Catheter 6 - Adhered |
| **Bs2** | *Bacillus subtilis* | Catheter 7 - Adhered |
| **Bs3** | *Bacillus subtilis* | Catheter 7 – Non Adhered |
| **Bs4** | *Bacillus subtilis* | Catheter 6 – Non Adhered |

Our data showed a high percentage of bacterial catheter colonization. Several authors have reported that the catheter colonization rate exceeds the urinary infection rate, indicating a significant inconsistency between urinary infections and catheter colonization. This discrepancy complicates the estimation of stent colonization (69, 70). Therefore, a thorough study should be carried out regarding the prevalence of the etiological agents and their antimicrobial susceptibility in double-J catheters, to improve treatments of long-term catheterized patients.

### Virulence related phenotypes

Measuring experimental phenotypic outcomes could predict the UP potential infectivity. Therefore, to understand the virulence strategies of the strains isolated here, we performed an extensive set of phenotypic microbiological assays: biofilm formation capacity, amyloid-type fiber, cellulose and mucoid substance production as components of the extracellular matrix, motility, siderophore production and hemolytic capacity, as well their ATB resistance profile. These phenotypes are summarized in **Table 2** and described below.

**Table 2.** Virulence associated-phenotypes.

|  |  | Biofilm formation |  |  |  |  |  | Amyloid-type fibers production | Cellulose production | Motility | Hemolytic capacity | Siderophores production | MDR phenotype |
| --- | --- | --- | --- | --- | --- | --- | --- | --- | --- | --- | --- | --- | --- |
|  |  | Culture conditions |  |  |  |  |  |  |  |  |  |  |  |
| UP isolate |  | M63 | LB | MacConkey | TSB | BHI | Urine | LB+CR+BB | LB+CW | 0.3%BHI | Blood agar | CAS medium | MH medium |
| *E. coli* | Ec1 | + | + | - | ++ | - | ++ | + | - | + | $\gamma$ | + | + |
| | Ec2 | ++ | - | - | +++ | + | ++ | + | - | + | $\alpha$ | + | + |
| | Ec3 | +++ | - | - | ++ | + | +++ | - | - | + | $\gamma$ | + | + |
| *K. pneumoniae* | Kp1 | +++ | ++ | +++ | +++ | ++ | +++ | - | + | - | $\gamma$ | - | + |
| | Kp2 | ++++ | + | ++++ | ++++ | + | ++++ | - | + | - | $\gamma$ | + | + |
| | Kp3 | ++++ | ++ | + | +++ | ++ | ++++ | - | + | - | $\gamma$ | - | + |
| | Kp4 | ++++ | + | ++++ | ++++ | + | ++++ | - | + | - | $\gamma$ | + | + |
| *S. aureus* | Sa1 | + | + | +++ | + | ++ | +++ | + | + | + | $\alpha$ | - | - |
| *S. epidermidis* | Se1 | - | + | - | + | + | +++ | + | + | ++ | $\alpha$ | - | + |
| | Se2 | - | + | - | +++ | +++ | +++ | + | + | ++ | $\alpha$ | - | + |
| | Se3 | - | - | - | - | ++ | +++ | + | + | ++ | $\gamma$ | - | + |
| | Se4 | - | - | +++ | +++ | ++ | +++ | + | + | - | $\alpha$ | + | - |
| | Se5 | ++ | + | ++++ | ++ | + | +++ | + | - | - | $\beta$ | - | - |
| | Se6 | - | | - | ++ | + | ++ | + | + | ++ | $\alpha$ | - | + |
| *E. faecalis* | **Ef1** | +++ | + | ++++ | +++ | ++ | +++ | - | + | + | $\alpha$ | - | + |
| | **Ef2** | +++ | - | +++ | +++ | +++ | +++ | - | + | + | $\alpha$ | - | + |
| | **Ef3** | ++ | - | ++ | +++ | ++ | +++ | - | + | ++ | $\alpha$ | - | + |
| | **Ef4** | +++ | - | + | +++ | +++ | +++ | - | + | + | $\alpha$ | - | + |
| | **Ef5** | +++ | + | +++ | ++ | ++ | +++ | - | + | + | $\alpha$ | - | + |
| | **Ef6** | +++ | + | ++ | ++ | +++ | +++ | - | + | - | $\gamma$ | - | + |
| *B. pumilus* | **Bp1** | - | + | - | ++ | ++ | +++ | + | + | + | $\beta$ | - | + |
| *B. megaterium* | **Bm1** | - | + | - | ++ | ++ | +++ | + | + | - | $\gamma$ | - | - |
| | **Bm2** | ++ | + | - | ++ | ++ | +++ | - | - | - | $\gamma$ | + | - |
| *B. subtilis* | **Bs1** | ++ | ++ | - | ++ | +++ | +++ | + | + | + | $\beta$ | + | - |
| | **Bs2** | ++ | ++ | - | - | +++ | +++ | + | + | + | $\beta$ | + | - |
| | **Bs3** | + | ++ | - | - | +++ | +++ | + | + | + | $\beta$ | + | - |
| | **Bs4** | + | ++ | - | - | +++ | +++ | + | + | + | $\alpha$ | + | - |

Phenotypes were represented as follow:
**Biofilm formation:** -, non-biofilm producers; +, weak biofilm producers; ++, moderate biofilm producers; +++, strong biofilm producers; and +++, robust biofilm producers.
**Amyloid-like fiber:** +, presence; -, absence.
**Cellulose production:** +, presence; -, absence.
**Siderophore production:** +, presence; -, absence
**Motility:** -, no motile; +, colony diameter between 0.5 and 1 cm; and ++, colony diameter more than 1 cm.
**Hemolytic capacity:** $\alpha$ , partial lysis; $\beta$ , complete lysis; $\gamma$ , no hemolysis.
**MDR phenotype:** +, MDR; -, No MDR

#### Biofilm formation

Biofilm formation capacity of all isolates was determined for up to 72 h using the crystal violet method. Cells were grown in M63, MacConkey, BHI, TSB and urine culture media for different times. According to the Stepanovic criteria the value of **A_595nm_C** was established for each condition as described in Material and Methods. Results in **Table 2** showed that the biofilm formation ability of the *E. coli*, *S. epidermidis* and *Bacillus* spp. strains varied depending on the employed culture media, being higher in TSB, BHI and urine medium. *S. aureus*, *E. faecalis* and most of the *K. peumoniae* isolates, were considered strong or robust biofilm formers in all tested media. Differences in the ability to form biofilms among genus and conditions are typical in clinical isolates (71–73). It is well known that biofilm formation is one of the most relevant virulence mechanisms used by UP during UTI (3, 71, 74). Indeed, it was reported that microorganisms living in these structures are even more virulent and resistant to ATB. Therefore, the ability of UP to form biofilms in urinary catheters and in the urothelium is a crucial factor for the persistence and recurrence of UTI (75). Also, the release of bacteria from biofilm to the bloodstream can lead to a widespread infection, especially in immune compromised patients (76–78). Description of biofilm formation capacity of the isolates carried out here, is not determining to characterize an isolate as no biofilm former, since in other circumstances, such as some clinical settings, they may have a greater capacity for adhesion and biofilm formation. Thus, subjecting the isolates to several experimental conditions such as the different media used in this work, can suggest their potential as biofilm former. In addition, to visualized biofilm structure, a qualitative analysis by confocal laser scanning microscopy (CLSM) and scanning electron microscopy (SEM) was carried out on 4 selected UP isolates in BHI medium (**Fig. S1 and S2**). Results showed that *S. aureus* Sa1, *E. coli* Ec1, *K.pneumoniae* Kp2 and *E. faecalis* Ef5 strains were able to adhere to the glass surface. Although surface coverage of Sa1 and Ec1 was relatively lower compared to Kp2, all these strains demonstrated a strong adhesion, presenting characteristic agglomerates. The biofilm surface of the Sa1, Ec1 and Kp2 strains showed a “Christmas tree forest” appearance with internal water channels. These structures, characterized by towering growth and interconnected channels, allow for efficient nutrient and waste exchange, enabling the biofilm to sustain itself over time (79). It is worth to mention that Ef5 isolate was not able to form an structured biofilm, thus, it was not possible to reconstruct the 3D structure. Additionally, when SEM was performed (**Fig. S2**), all isolates tested were also capable to adhere to the glass surface. However, the Sa1 isolate was not able to form a structured biofilm in this assay, where dispersed cells were observed. The typical coccus-shaped cells of Sa1 was observed, with a smooth appearance and less than 1 μm in size, which, in some fields, appeared in chains approximately with six cells long. In the Ec1 strain micrograph, a significant coverage of the glass surface was observed. Cells were approximately 1 μm long with a rough appearance. In addition, a delicate layer encapsulating groups of cells interspersed by channels was observed (indicated by an arrow in the insert), probably indicating the presence of extracellular matrix, as previously described for this genus (80). In the Ef5 isolate, a large number of cells clusters surrounded by abundant extracellular material were found. These cells exhibited the typical cocci shape with a size of less than 1 μm. Finally, the Kp2 isolate formed a strong biofilm, with protrusions that rise from the surface, as observed by CLSM. Pili, curli or nanotube-like structures were observed, with abundant extracellular matrix surrounding the bacteria (arrows in inserts of **Fig. S2**). These appendages have been reported to play essential roles during cell adhesion and biofilm formation (81, 82).

The discrepancies observed in the biofilm formation ability among the different isolates on polystyrene and glass surfaces were not surprising since surface topography is a parameter that significantly influences on microbial adhesion (83).

#### Colony morphotypes

Determination of morphology of microbial colonies and the ability to bind dyes can be used to describe, identify and characterize microorganisms, and serve as an indicator of the microbial physiology and metabolism. The binding to CR and Brilliant blue dyes and the interpretation of the results obtained is specific for each bacterial genus. **Table 2** and **Fig S3A** show phenotypes of bacterial colonies obtained in LB plates supplemented with CR and Brilliant Blue. When curli or other amyloid-like fibers were studied, it was observed that only *Escherichia*, *Staphylococcus* and *Bacillus* genus exhibited a colony morphotype characteristic of fiber or adhesion-type component production. Based on the morphotypes classification, *E. coli* isolates showed a *ras* phenotype (**Fig S3A**), indicating a moderate production of fimbria curli and cellulose. *S. epidermidis* isolates Se1, Se2, Se3 and Se6 showed clear, flat, circular colonies with rough edges at 24 h, turning into red, wrinkled, irregularly shaped at 96 h (**Fig S3A)**. According to the colony morphotype, *Staphylococcus* spp. isolates were considered as matrix producers (43). On the other hand, *Bacillus* isolates, Bp1, Bs1, Bs2, Bs3 and Bs4, showed dark/red colonies with central black halos, with rough, irregular, and lobed margins (**Fig S3A)**. This morphotype, indicates production of TasA proteins (45), knowing as the major amyloid-type protein of the biofilm matrix in *Bacillus* spp. (84–86). *K. pneumoniae* or *E. faecalis* strains were characterized as white colonies with pink/reddish centers and a mucoid appearance (**Fig S3B**). The distinctive mucoid morphology of *K. pneumoniae* isolates indicated a high capsule production (**Table 2**). This phenotype was reported as one of the main virulence factors of *K. pneumoniae* involved in adhesion to host cells and biofilm formation (87, 88). The presence of a pigmented ring stands out in Ef5 isolate was consistent with biofilm formation capacity (44).

Cellulose production is another relevant extracellular matrix component for biofilms and several virulence traits, since it provides an important physical barrier to antimicrobials (89–91). Besides the observed RC colonies morphotypes, the cellulose production was also visualized in LB-agar plates supplemented with Calcofluor White, as described in Materials and Methods. As shown in **Table 2**, except for *E. coli*, all bacterial genus produced cellulose, as evidenced by fluorescent colonies when were irradiated with UV light (**Fig. S3**). No fluorescence was observed in *E. coli* colonies after irradiation; only Ec3 isolate showed a slight fluorescence in the center of its colony, as expected for its *ras* morphotype.

Colony morphology depends on numerous environmental factors, such as media composition, temperature, and humidity, among others, which lead to the expression of specific genes or intercellular communication processes that control this process (Bokranz et al., 20(41, 42, 92, 93). Analysis of these components under a variety of conditions could determine the potential high capacity of the isolates to form a biofilm in UTI. Although correlation between morphology, expression of extracellular components and biofilm formation capacity in some bacterial isolates were denoted here (i.e. *B. subtilis* and *S. epidermidis* strains), it has not been generalized to all bacteria genera. For instance, it was observed that strong biofilm formers, such *K. pneumoniae* or *E. faecalis* strains, did not produce amyloid-type fibers (**Table 2).**

#### Bacterial motility

In some pathogenic bacteria, motility plays an essential role in the initial phase of infection (94), probably to overcome the electrostatic repulsion of cells and surfaces (95, 96). In addition, motility contributes to the colonization of different environmental niches by facilitating the spread of the infectious agent, crucial in the context of UTI (94). As observed in **Table 2**, all isolates showed and important motility capacity, except for *K. pneumoniae*, *S. epidermidis* Se4, Se5, *B. megaterium* and *E. faecalis* Ef6 strains. Although both *S. aureus* and *S. epidermidis* lack flagella, therefore are considered as non-motile bacteria, it was observed that Sa1, Se1, Se2 and Se3 isolates exhibited significant motility. Recent studies have reported flagellum-independent forms of motility in microorganisms of this genus (97). For instance, spreading involves a sliding movement, where bacteria spread radially outward from an inoculation site, forming multiple layers of densely packed cells (98, 99). *E. faecalis* is also regarded as a non-motile bacterium, due to the absence of flagella. However, in this study, most *E. faecalis* isolates showed motility, with the exception of the Ef6 strain. It was demonstrated an *E. faecalis* migration mechanism, where the synthesis and secretion of extracellular polysaccharides were required (100). Therefore, when cells are grown on semisolid agar, they penetrate and invade the medium, creating a “colony impression” (100). This mechanism is relevant for translocation through monolayers of human epithelial cells, conferring adaptive advantages during infection, since they would be able to translocate from the urinary tract to the bloodstream to colonize distant anatomical sites (101–103). In general, it was described that environmental isolates of *B. subtilis* exhibit a robust motility (104), in agreement with the exacerbated motility showed by the strains used in this work. There is a complex relationship between motility and biofilm formation, and depends on the environmental conditions or the bacteria requirements. A study conducted on *B. cereus* strains reported that bacterial motility influenced biofilm formation through 3 mechanisms: 1-motility is necessary for the bacteria to reach adequate surfaces to form biofilm; 2-motility promotes the recruitment of planktonic cells to invade the preformed biofilm; and 3-motility is involved in biofilm spreading and propagation (105). In relation to *B. subtilis* motility observed here, could be assumed that motile bacteria should become non-motile when transitioning to the biofilm state (72, 106).

#### Siderophores production

Iron, a vital nutrient required for bacterial growth, is highly restricted within human hosts (107, 108). Most UP strains encode several iron acquisition systems, such as siderophores to acquire iron sequestered by the host (109). Here, we tested siderophore production by the studied clinical isolates using the chrome azurol S (CAS) assay, described previously. All *E. coli* and *B. subtilis*, Kp2 and Kp4, Se4 and Bm1 strains were able to produce siderophores (**Table 2** and **Fig. S4A**). It was reported that siderophores production by different UP is an important characteristic that contributes to the potential virulence of these isolates, relevant in the CAUTI context (110–112).

#### Hemolytic capacity

Hemolysins are proteins secreted by bacteria that constitute another important virulence factor in UTI. These toxins can disrupt host cell signaling cascades, altering the inflammatory response and inducing cell death (113, 114). It is well known that α-hemolysin, HlyA, can stimulate the epithelial barrier rupture, producing the translocation of the bacteria from the urinary tract to the bloodstream (115). **Table 2** shows that 19.2% of the isolates exhibited total hemolysis (β hemolysis) with defined lysis zones (**Fig. S4B**). A partial hemolysis (α hemolysis) was carried out by 42.3% of the isolates, while the remaining 38.5% did not present hemolytic capacity. As observed in **Table 2**, *Staphylococcus* strains exhibited partial or total hemolysis capacity. It is well known that hemolytic capacity is one of the most common virulence factors of coagulase-positive (*S. aureus*) and coagulase-negative (*S. epidermidis*) staphylococci (116). In *E. faecalis* isolates, only a partial hemolysis was observed, as expected, since several studies described that this genus is not able to produce total hemolysis (44, 117).It is worth to mention that 4 out of the 5 isolates exhibiting total hemolysis belong to the *Bacillus* spp. genus. Similar results were obtained with *Bacillus* strains isolated from river water samples with fecal contamination, where all isolates presented total hemolytic capacity (118). The pathogenicity of *Bacillus* spp. has been poorly investigated, except for *B. cereus*, a known pathogen, whose pathogenic potential has been related to the secretion of several virulence proteins, such as hemolysins phospholipases, cytotoxin K (CytK) and proteases (119, 120), and to diverse motility factors, such as swimming and swarming (121).

Taken together, the ability to form biofilms, produce extracellular components like capsules and cellulose, along with virulence factors such as hemolysins and siderophores, are critical elements in the pathogenicity of UP associated with CAUTI. These factors not only facilitate adhesion and colonization of devices, but also confer resistance to antimicrobial treatments, complicating the resolution of infection.

### Antibiotic susceptibility profile

Knowledge of antibiotic susceptibility (AS) pattern of UP is necessary to characterize the microorganisms and achieve effective management of UTI in the double-J stent context. Therefore, antibiograms using conventional ATB were performed for each isolate (**Table S2**). Results in **Table 2** show isolates that presented a MDR phenotype. *E. coli* isolates (Ec1, Ec2, and Ec3) were MDR, being resistant to 6 out of 12 ATB tested. As observed here, several authors described that most UPEC isolates were resistant to first-generation cephalosporins and fluoroquinolones (122–126). Also, IMI susceptibility suggests that carbapenems still offer a viable treatment option for MDR *E. coli* infections, although the emergence of carbapenem-resistant strains is an ongoing concern (127). *K. pneumoniae* isolates also exhibited a MDR profile, being susceptible only to IMI and AK (**Table S2**), as previously describes by other authors (128). *K. pneumoniae* has been highlighted as a major pathogen involved in outbreaks of healthcare-associated infections. Among the pathogens responsible of UTI, both *E. coli* and *K. pneumoniae* are of particular concern, since they have acquired plasmids encoding extended-spectrum β-lactamases (ESBLs) (129–133). These plasmids rapidly propagate resistance to third-generation cephalosporins, as well as other ATB, thus increasing prevalence and diffusion of ESBL microorganisms (131). Although in this work we did not analyzed whether the isolates were ESBL producers, resistance to second and third generation cephalosporins was observed in *K. pneumoniae* isolates (**Table S2**). Regarding to *Staphylococcus* genus, only Se1, Se2, Se3 and Se6, show a MDR phenotype (**Table 2**). Multi-drug resistant *S. epidermidis* is increasingly recognized as a cause of opportunistic infections, particularly in patients with indwelling medical devices (134). These infections can be difficult to treat due to biofilm formation and the ability of *S. epidermidis* to acquire resistance genes through horizontal gene transfer. Our results are in agreement with those reported by Socohou et al. (2020), where *S. epidermidis* strains isolated from surfaces of medical materials exhibited high susceptibility to gentamicin and ciprofloxacin, whereas S*. aureus* isolates showed greater sensitivity to chloramphenicol and vancomycin (135). Regarding to *E. faecalis* strains, all isolates showed a MDR profile. The observed resistance to ERY, CLI, OXA and FOX (**Table S2**) was particularly alarming. Additionally, vancomycin-resistant enterococci (VRE) strains were identified, as indicated by their VAN and ERY resistance (Ef5 and Ef6 strains). VRE appearance, particularly in catheterized patients, is a growing problem worldwide, with significant clinical implications due to the limited treatment options available (136–139). *E. faecalis* resistance to multiple ATB, such as ERY, CLI, OXA, and FOX, represents a significant clinical concern, particularly in hospital and healthcare settings. *E. faecalis* resistance to macrolides, such as erythromycin, and lincosamides, like clindamycin, is important since these ATB are frequently used as an alternative to β-lactams in patients with penicillin allergies. The ability of *E. faecalis* to survive in the presence of these ATB can lead to persistent infections, especially in vulnerable patients undergoing surgeries, catheter placements, or long-term antibiotic therapy (136). Additionally, the spread of these resistant strains can compromise infection control efforts in hospitals, leading to outbreaks of difficult-to-treat infections. Results show that *Bacillus* spp. presented limited resistance to the tested ATB. Particularly, only Bp1 strain exhibited resistance to several antimicrobials, such as CAZ, ERY, CLI, OXA y FOX, while Bm1 and Bs3 strains were resistant only to AMP. Similar results for ATB resistance in *B. pumilus* and *B. subtilis* were informed by Adamski et al. 2023 (140). Also, since most *Bacillus* species were not considered human pathogens, but simply saprophytic skin microorganisms, no antimicrobial susceptibility studies are usually performed for this genus (55, 65). As environmental organisms, *Bacillus* are not frequently subjected to the selective pressures found in hospital settings, where the continuous use of ATB often leads to the development of resistance (141). This limited exposure may contribute to the observed lower rates of resistance compared to other pathogens commonly encountered in healthcare environments.

### Presence of genes that encodes for virulence factors

The ability of UP isolates to cause different diseases is mediated by multiple virulence factors, including the expression of genes associated with the production of adhesins, toxins, siderophores, and secretion systems, among others (21, 22). These factors are involved in the colonization of specific host surfaces, evasion of immune defenses, or direct damage to cells and tissues (22). *E. coli* isolates showed a consistent pattern in virulence gene profiles, containing genes for iron acquisition systems (*iutA* and *iroN*), and the hemolysin-encoding gene *hlyA*. Additionally, the Pap fimbriae (*papA*) and cytotoxic necrotizing factor (*cnf-1*) genes were also detected in all isolates, suggesting a highly virulent phenotype. *iutA* and *iroN* are known to enhance bacterial survival by allowing iron acquisition in iron-limited environments, such as the human body (142). Kanamaru et al. (2003) reported the prevalence of putative uropathogenic virulence factors in 427 *E. coli* strains and reported a significant prevalence rate of the *iroN* gene in the UTI isolates (143). The presence of *hlyA* and *cnf-1* further suggests that these isolates have the potential to cause severe infections, as both genes are linked to tissue damage and inflammatory responses, as usually observed in UPEC (144, 145). Soto et al. (2007) reported that *papA* and *hlyA* genes were present in 42% and 36% of the UPEC strains analyzed, respectively (146). It is worth to mention that 50% of UPEC produce α-hemolysin, and its expression is strongly associated with symptomatic UTI (4).

All *K. pneumoniae* isolates were positive for all virulence genes tested (**Table 3**), indicating their potential to cause infections by adhering to host tissues and evading the immune system. *OmpK36* expression has been linked to ATB resistance (147), consistent with the MDR profile previously described for this genus (**Table 2** and **Table S2**). Also, the combination of adhesion factors and hemolysin suggests that these strains may be particularly proficient at establishing infections, particularly in immunocompromised patients (148).

**Table 4.** Presence of virulence factor coding genes.

| UP isolate |  | Genes |  |  |  |  |  |  |
| --- | --- | --- | --- | --- | --- | --- | --- | --- |
|  |  | *fimA* | *fimB* | *iutA* | *iroN* | *papA* | *cnf-1* | *hlyA* |
| *E. coli* | Ec1 | - | + | + | + | + | + | + |
|  | Ec2 | - | + | + | + | + | + | + |
|  | Ec3 | - | + | + | + | + | + | + |
|  |  | *fimA* | *entB* | *mrkD* | *ompK36* | *hlyA* |  |  |
| *K. pneumoniae* | Kp1 | + | + | - | + | + |  |  |
|  | Kp2 | + | + | - | + | + |  |  |
|  | Kp3 | + | + | - | + | + |  |  |
|  | Kp4 | + | + | - | + | + |  |  |
|  |  | *sea* | *ica* | *agrB* | *argC* | *geH* | *sdrG* |  |
| *S. aureus* | Sa1 | - | - | - | + | + | + |  |
| *S. epidermidis* | Se1 | - | - | - | + | + | + |  |
|  | Se2 | - | - | - | + | + | + |  |
|  | Se3 | - | - | - | + | + | + |  |
|  | Se4 | - | - | - | - | + | + |  |
|  | Se5 | - | - | - | - | + | + |  |
|  | Se6 | - | - | - | + | + | + |  |
|  |  | *gelE* | *srt* | *efaA* | *ace* | *cylA* |  |  |
| *E. faecalis* | Ef1 | + | - | + | + | + |  |  |
|  | Ef2 | + | - | + | + | + |  |  |
|  | Ef3 | + | - | + | + | + |  |  |
|  | Ef4 | + | - | + | + | + |  |  |
|  | Ef5 | + | - | + | + | + |  |  |
|  | Ef6 | - | - | - | - | - |  |  |
|  |  | *entA* | *entB* | *cytK* | *clpC* | *sdrG* | *hlyA* |  |
| *B. pumilus* | Bp1 | - | - | - | - | - | - |  |
| *B. megaterium* | Bm1 | - | - | - | - | - | - |  |
|  | Bm2 | - | - | - | - | - | - |  |
| *B. subtilis* | Bs1 | + | + | - | + | + | + |  |
|  | Bs2 | - | - | - | - | - | - |  |
|  | Bs3 | - | - | - | - | - | - |  |
|  | Bs4 | + | + | - | + | + | + |  |

The *S. aureus* isolate presented virulence genes related to the accessory gene regulator (agr) system, such as *argC*, and *geH*. This system is involved in the expression of toxins and surface proteins that aid to immune evasion and tissue invasion (149). The agr system is well-documented as a key virulence regulator in *S. aureus*, controlling the production of toxins, such as hemolysins and proteases (149). Additionally, the *sdrG* gene, associated with fibrinogen-binding proteins that contribute to biofilm formation and immune evasion, was observed in this isolate. The presence of this gene indicates an enhanced biofilm-forming capacity, exhibiting a strong resistance to immune clearance and ATB treatment (150). *S. epidermidis* isolates also contained the *agrC*, *geH* and *sdrG* genes, all associated with biofilm formation (151). This ability enhances the survival of *S. epidermidis* on indwelling medical devices, such as catheters and prosthetic joints (134). The presence of these virulence factors suggests that these *S. epidermidis* isolates could be well-adapted and cause persistent infections in healthcare settings.

Except for Ef6, all *E. faecalis* isolates were positive for *gelE*, *efaA*, *ace* and *cylA* genes. The presence of these virulence factors highlights the pathogenic potential of *E. faecalis*. Gelatinase, encoded by *gelE* gene, is a protease that degrades host tissues and promotes biofilm development by facilitating bacterial adhesion (152). The Ace protein is involved in adhesion to extracellular matrix proteins, and is important for the host tissues colonization and the establishment of biofilms on medical devices (153). EfaA is also involved in bacterial adherence and is considered essential for biofilm formation during infection (154). The expression of these genes are consistent with the considerable biofilm formation observed in these isolates. Finally, cytolysin (*cylA*) is involved in the lysis of red and white blood cells, contributing to immune evasion and tissue destruction (155). This result is in agreement with the partial hemolytic capacity exhibited by these isolates.

Among the analyzed *Bacillus* species, only *B. subtilis* (Bs1 and Bs4) exhibited *hlyA* and *sdrG* genes. This was unexpected, since *Bacillus* spp. are generally regarded as environmental bacteria with limited clinical relevance (156). However, recent studies by Bianco et al. (2021) and Fayanju et al. (2024) pinpointed specific virulence genes in *B. subtilis* strains that contribute to their pathogenicity (157, 158). In addition to virulence factors, the antimicrobial resistance profiles of Bacillus strains play a critical role in determining their pathogenicity and potential impact on CAUTI contexts.

### Assessment of stress response phenotypes

In the urinary tract, bacteria are exposed to a variety of host-mediated stress responses, such as osmotic stress, pH changes, reactive oxygen species (ROS) from the immune system, and nutrient limitation (159). Therefore, pathogens develop adaptive advantages to cope with these environmental stressors.

All isolates were exposed to pH 7, 5, and 2.5 for 2 h to cover the expected pH that bacteria may encounter in the gut, urine, or during neutrophil attack (**Fig. 1**). Results showed that both, Gram-positive and Gram-negative isolates significantly decreased their viability when exposed to pH 2.5. Only Ec2, Sa1, Bs1 and Bs4 strains were tolerant to all tested pH conditions (**Fig. 1A**). The decreased viability when exposed to pH 2.5 is consistent with previous findings that show that acidic conditions can severely compromise bacterial membrane integrity and metabolic function in both bacterial genera (160, 161).

**Figure 1.**
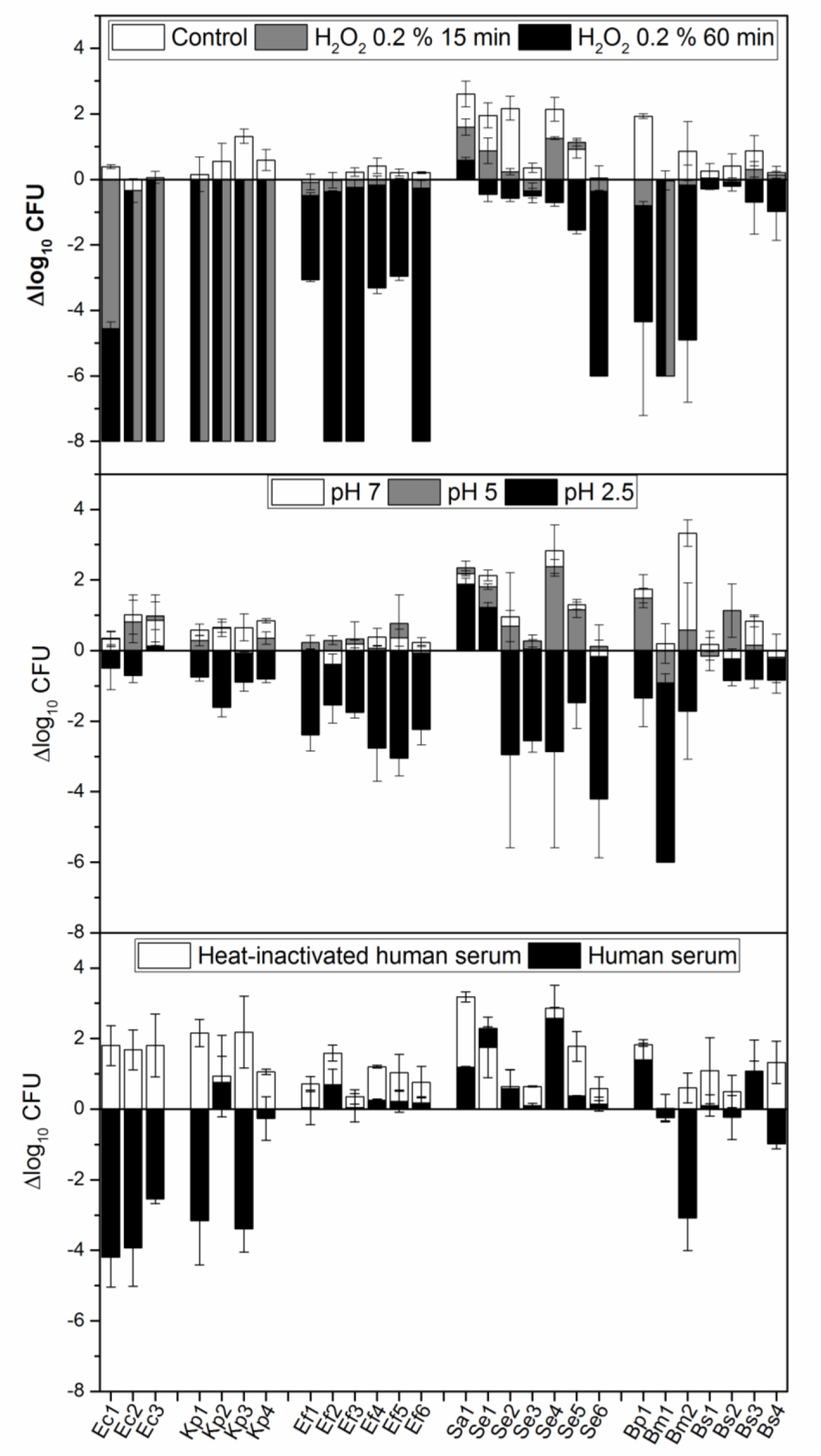
pH, H_2_O_2_, and human serum tolerance assays. (A) A total of 10^8^ CFU/mL of each isolate were inoculated into BHI containing 0.2% H_2_O_2_ and maintained statically at room temperature. Samples were taken and serially diluted at 0, 15, 60 min. (B) From an overnight culture, UPEC isolates were diluted to 10^8^ CFU/mL in BHI buffered to pH 7 (control), 5, or 2.5, as indicated. Cultures were incubated for 2 h at 37°C with aeration. (C) Isolates were cultured overnight in BHI and washed and resuspended in PBS, and then 10^8^ CFU/mL was added to 100% complete human serum and heat-inactivated human serum. Samples were incubated statically at 37°C for 60 min. After incubations, CFU/mL was determined. Data represent the mean ± SD of at least four independent experiments. One-way ANOVA was performed with Fisher test with a p-value of 0.05.

In order to study oxidative stress, survival of the tested isolates exposed to H_2_O_2_ was assessed. *E. coli* and *K. pneumoniae* strains (both Gram-negative bacteria) were highly sensitive to 0.2 % H_2_O_2_ even after 15 min (**Fig. 1B**). However, Gram-positive bacteria were more resistant to the mentioned stress, displaying variable susceptibility when exposed for up to 1 h (**Fig. 1B**). The higher resistance in Gram-positive isolates could be due to differences in their cell wall structures and antioxidant defense systems (162).

Complement-mediated killing is also an important innate immune defense that can be tested by measuring bacterial survival in human serum. Only *E. coli* isolates, Kp1, Kp3 and Se5 strains were partially susceptible to serum (**Fig. 1C**). This is in agreement with the fact that, although the immune pressure of serum complement, all virulent pathogens capable of inducing active infections have evolved immune evasive strategies that primarily target the complement system (163).

The significant resistance to exogenous stressors observed in many isolates implies that these bacteria may have a high survival rate during infections, potentially leading to chronic or recurrent infection.

### Principal Component Analysis of Phenotypic Diversity

The Principal Components Analysis (PCA) plot provides a visual summary of the relationships between five different bacterial genera (color- and shape-coded by taxonomic group) based on eleven phenotypic traits: antimicrobial resistance, biofilm formation, hemolytic capacity, motility, siderophores production, virulence factor coding genes, curli production, cellulose production and H_2_O_2_, pH 2.5, and human serum resistances (**Fig. 2**). The variable “pH 5 resistance” was excluded from the PCA since all bacteria were resistance (constant value). The X-axis (PC1) explained 27% of the variance, while the Y-axis (PC2) explained 23%, together accounting for 50% of the total variance, which allowed an adequate interpretation of the characteristics associated with virulence (**Fig. 2**).

**Figure 2.**
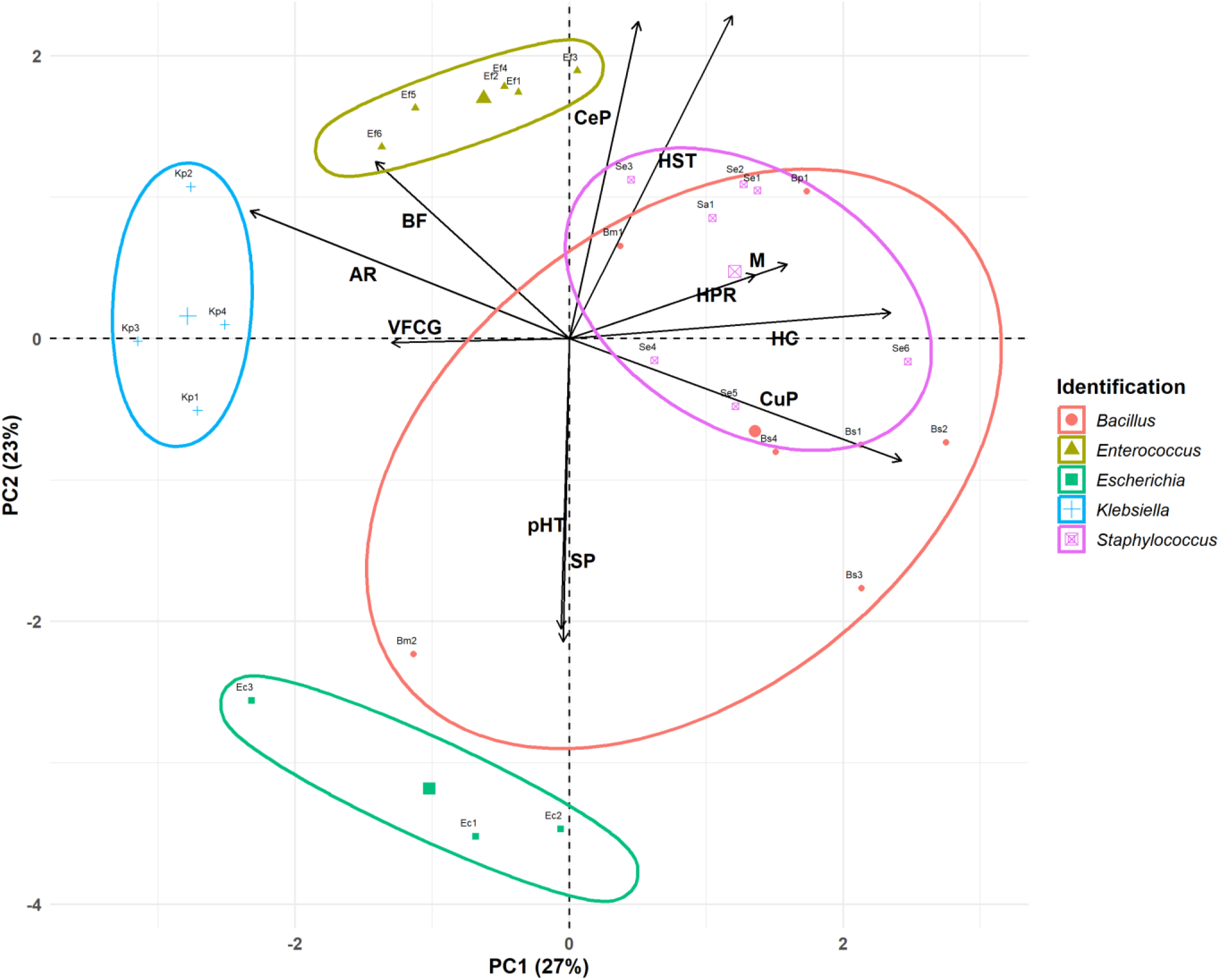
**Principal Components Analysis,** including antimicrobial resistance (AR), biofilm formation (BF), hemolytic capacity (HC), motility (M), siderophores production (SP), virulence factor coding genes (VFCG), curli production (CuP), cellulose production (CeP), H_2_O_2_ resistance (HPR), pH 2.5 resistance (pHR), and human serum resistance (HSR).

Based on PCA loadings, PC1 primarily distinguished bacteria by traits such as antimicrobial resistance (-0.45), biofilm formation (-0.27), virulence factor coding genes (-0.25) on the left side, versus curli production (0.47), hemolytic capacity (0.46), motility (0.31), and H_2_O_2_ resistance (0.26) on the right side. PC2 differentiates bacteria based on human serum resistance (0.48), cellulose production (0.47), and biofilm formation (0.27) (upper quadrant) versus siderophores production (-0.45) and pH 2.5 resistance (-0.43) (lower quadrant). It is important to highlight that biofilm formation contributes equally to both components of the PCA. Both components may reflect a balance between pathogenic potential and resilience against host defenses.

Each bacterial genus was enclosed within shaded ellipses, representing the clustering tendency for each bacterial group. The within-cluster sum of squares (WSS) for each group was calculated to reflect the internal variability of the clusters formed by the algorithm.

PC1 segregated the bacteria along the x-axis, positioning the clusters of *Klebsiella* (WSS: 1.53), *Enterococcus* (WSS: 1.56), and *Escherichia* (WSS: 3.12) to the left. These clusters were associated with antimicrobial resistance and the presence of virulence factor-coding genes. The distinction among these genera was marked by PC2, which separated the bacteria along the y-axis. *Enterococcus* and *Klebsiella* were located at the top of the graph, linked to biofilm formation and cellulose production. In contrast, *Escherichia* was positioned at the bottom of the graph associated with siderophore production and resistance to pH 2.5. Our results confirmed that *Klebsiella* and *Enterococcus* isolates, with robust and strong biofilm-forming abilities, respectively, were resistant to multiple antimicrobials, suggesting a direct relationship between antimicrobial resistance and biofilm formation capacity (see **Tables 1** and **3**). Although the UP isolates were derived from patients who had not received antibiotic therapy in the 10 days prior to the device removal procedure, it is important to consider the potential impact of previous extended antimicrobial treatments, which could have contributed to the observed resistance. Antibiotic therapy is crucial for the treatment of UTI, but the rising prevalence of MDR UP in recent years represented a significant challenge for their effective management (164). On the other hand, the combination of increased motility with a high prevalence of virulence factors observed in *Enterococcus* strains suggested that these isolates may be particularly difficult to treat, becoming a significant risk in clinical settings.

*Staphylococcus* (WSS: 2.84) was clustered in the right center of the graph. According to PC1, this genus was linked to hemolytic capacity, motility, and curli production. Regarding PC2, this cluster was associated with cellulose production, human serum resistance, and biofilm formation.

The most heterogeneous group was *Bacillus* (WSS: 3.99), with members distributed across three quadrants of the biplot. Taking PC1 as a reference, all members were associated with curli production (except Bm2). Regarding PC2, all members were clustered by cellulose production (except Bm2) and human serum resistance (except Bm2 and Bs3). *B. subtilis* were linked in the bottom right, associated to siderophores production and pH 2.5 resistance (PC2). Also, they were related to peroxide resistance, motility, and total hemolytic capacity. Due to the *Bacillus* cluster’s variability, one characteristic that cannot be fully appreciated was the strong biofilm formation, as all bacteria of this genus exhibit this trait. This ability could also allow it to persist in medical or industrial environments, though it is rarely associated with human infections. High curli production and strong biofilm formation observed in both *Staphylococcus* and *Bacillus* suggest a robust capacity for adhesion and colonization on both abiotic and biotic surfaces. These microbial attributes may have implications for their virulence, increasing the risk of systemic infections.

### Co-isolates pair interaction

Polymicrobial interactions can modify the pathogenic potential of one organism over the others. Therefore, microbes co-existing in complex communities must employ diverse mechanisms, such as cross-feeding, cooperation, competition, and immune modulation, to not only shape the composition of the bacterial community, but also influence interactions with the host, affecting the progression from colonization to infection (165). Although an exhaustive characterization of each isolate grown as a pure culture was carried out, it is essential to study their behavior in polymicrobial systems. As mentioned before, in 5 of the 8 catheters, more than one bacterial species was isolated. Notably, *Bacillus* spp. were present in 4 catheters, which allowed us to study their role in the interaction with microorganisms typically regarded as pathogens. Therefore, we investigate the interaction between four co-isolated pairs obtained simultaneously from the same catheter: Ef5/Bp1, Kp2/Bm2, Se3/Bs1, and Se4/Bs2. Firstly, we performed interaction assays in solid medium, with deferred and simultaneous inoculation of the isolates. As shown in **Fig. 3A**, Bp1 inhibits the growth of Ef5 when inoculated simultaneously and when it was inoculated first in deferred assays. However, when Ef5 was firstly inoculated, a change in the colony complexity was observed in the interaction zone, denoted by the loss of the characteristic wrinkling of the Bp1 colony. Regarding Kp2/Bm2, Kp2 induced a slight transparency in the Bm2 colony in the interaction zone, which can be interpreted as an inhibition of Bm2 when both bacteria grew simultaneously. This phenotype was exacerbated when Kp2 growth first in the deferred assay (**Fig. 3A**). When *S. epidermidis*/*B. subtilis* interactions were analyzed, results show that, in both cases, *S. epidermidis* exacerbated the *B. subtilis* motility when bacteria grew simultaneously. Notably, this effect was more robust in the Se4/Bs2 pair. Also, an increase in the number of Se3 colonies associated with a greater induction of Bs2 motility. When *B. subtilis* Bs1 and Bs2 grown first, a significantly inhibition of their counterpart (Se3 and Se4, respectively) was observed (**Fig. 3A**). Behaviors between these bacteria were not surprising since it was recently described that *B. subtilis* actively responds to the presence of *S. epidermidis* in its proximity by two strategies: antimicrobial production and development of a subpopulation with migratory response (166). However, in this work, these behaviors depend on the moment in which the interaction occurs. Controls of single colonies growth at 24 and 48 h are shown in **Fig. S5.** Interspecie interactions vary significantly depending on whether the bacterial strains grow individually or share the same growing location. When some isolates grow first (e.g., Ef5, Bp1, and Bs1) the potential production of metabolites, including antimicrobial compounds or other agents that alter the local environment, may lead to the observed inhibition of the counterpart upon its arrival. However, this potential metabolite production may be modified in the presence of the other strain when both grow simultaneously, resulting in a different type of interaction. This observation raises the possibility that colonization order influences physiological adaptations and virulence expression in each species, potentially affecting infection progression and severity in the host.

**Figure 3.**
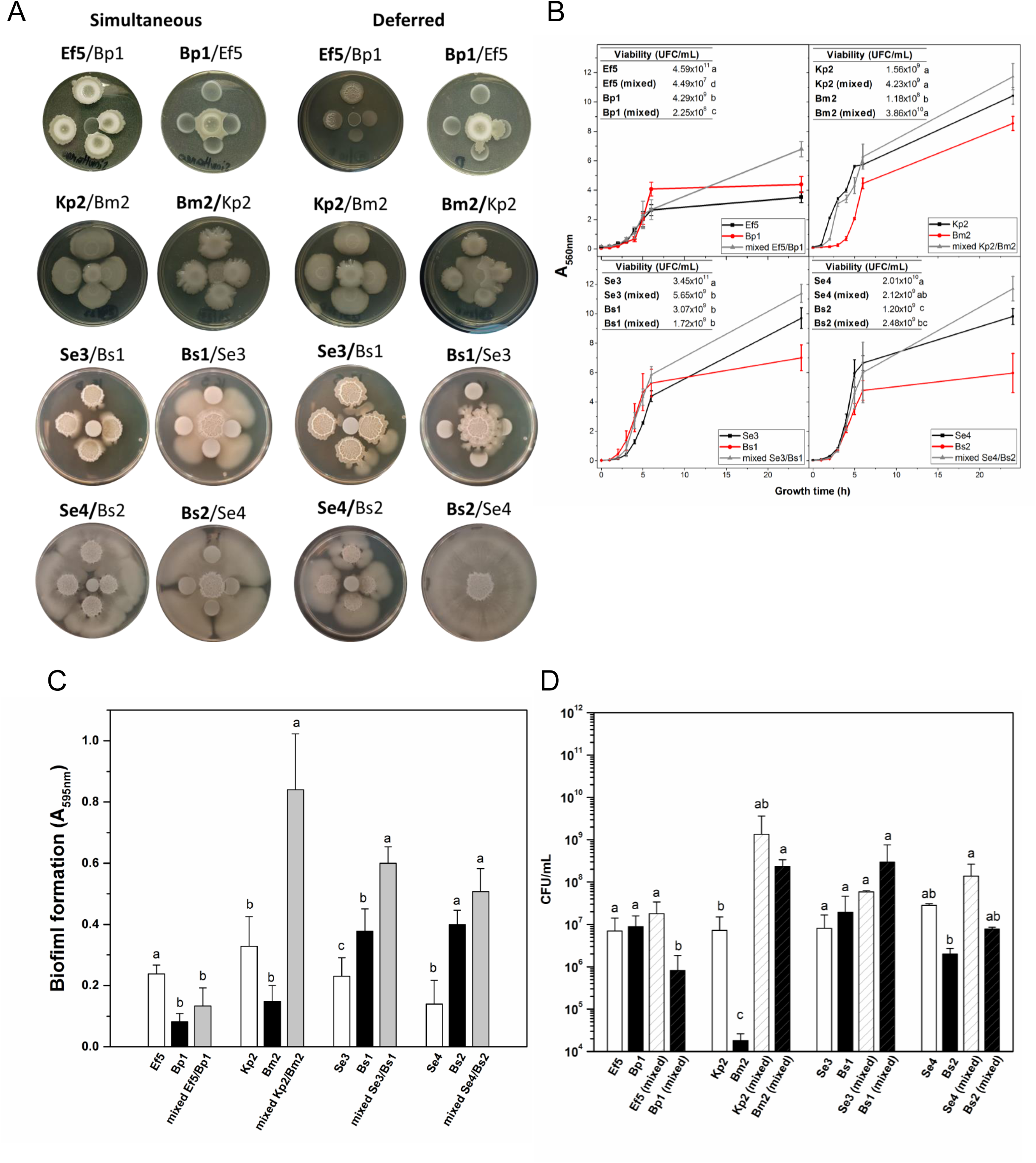
Interaction of co-isolated pairs. **(A)** In colony interaction assay aliquots of the co-isolated pair were inoculated at the same time (simultaneous) or 24 h post growth at 30°C (deferred) in BHI-agar plates. Bold letters in the name pair indicates de central bacterium. **(B)** Growth curves (A_560 nm_) of the axenic or mixed (Ef5/Bp1, Kp2/Bm2, Se3/Bs1, and Se4/Bs2) were carried out in BHI with agitation during 24 h. Inset: Isolate viability in axenic or mixed condition. **(C)** Biofilm formation (A_595nm_) and **(D)** cell viability (CFU/mL) was determined for the four co-isolated pair after 24 h of growth. Data represent the mean ± SD of at least four independent experiments. For each co-pairs interaction assay, different letters indicate significant differences according to One-way ANOVA performed with Fisher test with a p-value of 0.05.

Since bacteria in clinical settings are generally found as mixed cultures or communities that share a common environment, we decided to perform co-culture assays in agitated and static liquid media. Growth curves (OD) and viable cell counts (CFU ml^-1^) for both mixed and monocultures are shown in **Fig. 3B**. In general, mixed cultures in all pairs, except for the Kp2/Bm2, displayed higher OD values than monocultures after 24 h (*p*-value < 0.05). However, different results were observed when viable cell quantification was assessed. In mixed culture, Ef5 and Bp1 exhibited a lower viability compared to their respective monocultures (see inset table). In the Se3/Bs1 and Se4/Bs2 pairs, viable cells of *S. epidermidis* strains in mixed culture decreased by at least one magnitude order, suggesting that *B. subtilis* may exert an inhibitory effect on its growth. This inhibition may be attributed to the production of secondary metabolites by *B. subtilis*, such as surfactins or antimicrobial peptides, previously reported as antimicrobial substances (166, 167). On the other hand, in the Kp2/Bm2 pair, Kp2 viability remained unaffected when grown in mixed culture after 24 h, while Bm2 CFU/ml^-1^ increased in mixed cultures.

Considering that co-isolated pairs were attached to the catheters, biofilm formation of the four co-isolated pairs was evaluated on polystyrene plates after 24 h in mixed and axenic cultures (**Fig. 3C**). Results showed that biofilm formation was enhanced in three of four co-isolate pairs (Kp2/Bm2, Se3/Bs1, and Se4/Bs2), compared to single-species biofilms (**Fig. 3C**, left panel). To assess the contribution of each isolate to biofilm formation, bacterial viability was determined in the biofilms (**Fig. 3C**, right panel). As expected, in the co-cultures, where increased biofilm formation was observed, the number of viable cells was higher than in the monocultures. Different studies demonstrated that *B. subtilis* often facilitates biofilm development by producing extracellular polymeric substances (EPS), which provide structural support and protection to cohabitating species within the biofilm matrix (168). Regarding the Ef5/Bp1 pair, Ef5 viability remained similar in mixed and single-species biofilms, while the number of viable Bp1 cells was reduced in the mixed culture compared to the monoculture. This may be due to the production of antimicrobial peptides since *Enterococcus* spp. are one of the most frequent producers of bacteriocins (169). Our results showed that interactions that occurred between the different co-isolated pairs, reflected a combination of competitive, inhibitory, and cooperative behaviors in biofilm formation, consistent with the description of microbial cohabitation in this life-style (170).

The role of *Bacillus* spp. in these dynamics, particularly its ability to enhance biofilm formation and to inhibit certain pathogens, highlights the complexity of polymicrobial interactions in infection environments like CAUTI. The order of colonization may not only determine which microorganism dominates the niche, but may also have a direct impact on the clinical outcome for the patient. If a virulent bacterium colonizes first, it may alter the environment, either promoting or limiting subsequent colonization by other species, thereby influencing the overall virulence of the infection. Conversely, simultaneous colonization could lead to complex interspecies interactions with potentially unpredictable consequences for infection progression and treatment efficacy. These findings underscore the need to investigate the role of polymicrobial colonization in clinical infections and its impact on host response and antimicrobial treatment efficacy.

## CONCLUSION

The characterization of phenotypic and genotypic traits of UP isolates obtained from double-J catheters enhances the understanding of the potential virulence of these bacteria in clinical settings, particularly in the context of CAUTI. The observed virulence factors and interactions of *Bacillus* spp. suggest that this genus may act as an opportunistic pathogen, highlighting its potential clinical relevance in co-infections of the urinary tract. Overall, this work lays a foundation for advancing the knowledge of bacterial pathogenesis associated with CAUTI, offering crucial insights for effective clinical management and treatment strategies in cases of multi-bacterial urinary infections.

## ETHICAL CONSIDERATIONS

In the present study, no direct contact was maintained with the patient or his data. Thus, informed consent was not required.

## COMPETING INTERESTS

The authors declare that they have no competing interests.

## AUTHORS’ CONTRIBUTIONS

MGP, JVF and JMV designed the study. JVF, EC and JAE carried out virulence associated-phenotypes and antimicrobial profiles assays. CV performed identification of clinical isolates. JVF, JAE and JMV carried out stress tolerance assay. JVF and JAE executed interaction of co-isolated pairs experiments. EMH and LS performed virulence genes determination assay. JDA performed PCA analysis and general statistical analysis. MGP, JVF and JMV prepared the manuscript and participated in the analysis of data. VAR discuss and revised the manuscript for intellectual content. MGP, JMV and VAR contributed with reagents, materials and analysis tools. MGP, JVF and JMV wrote the paper. All authors read and approved the final manuscript.

## ACKNOWLEDGMENT

We gratefully acknowledge Dr. Martin Lescano for providing the catheters. This research was supported by Argentinean grants of the Consejo Nacional de Investigaciones Científicas y Técnicas (CONICET) and the Agencia Nacional de Promoción Científica y Técnica (ANPCyT).

## Supplementary Material

**Table S1.**
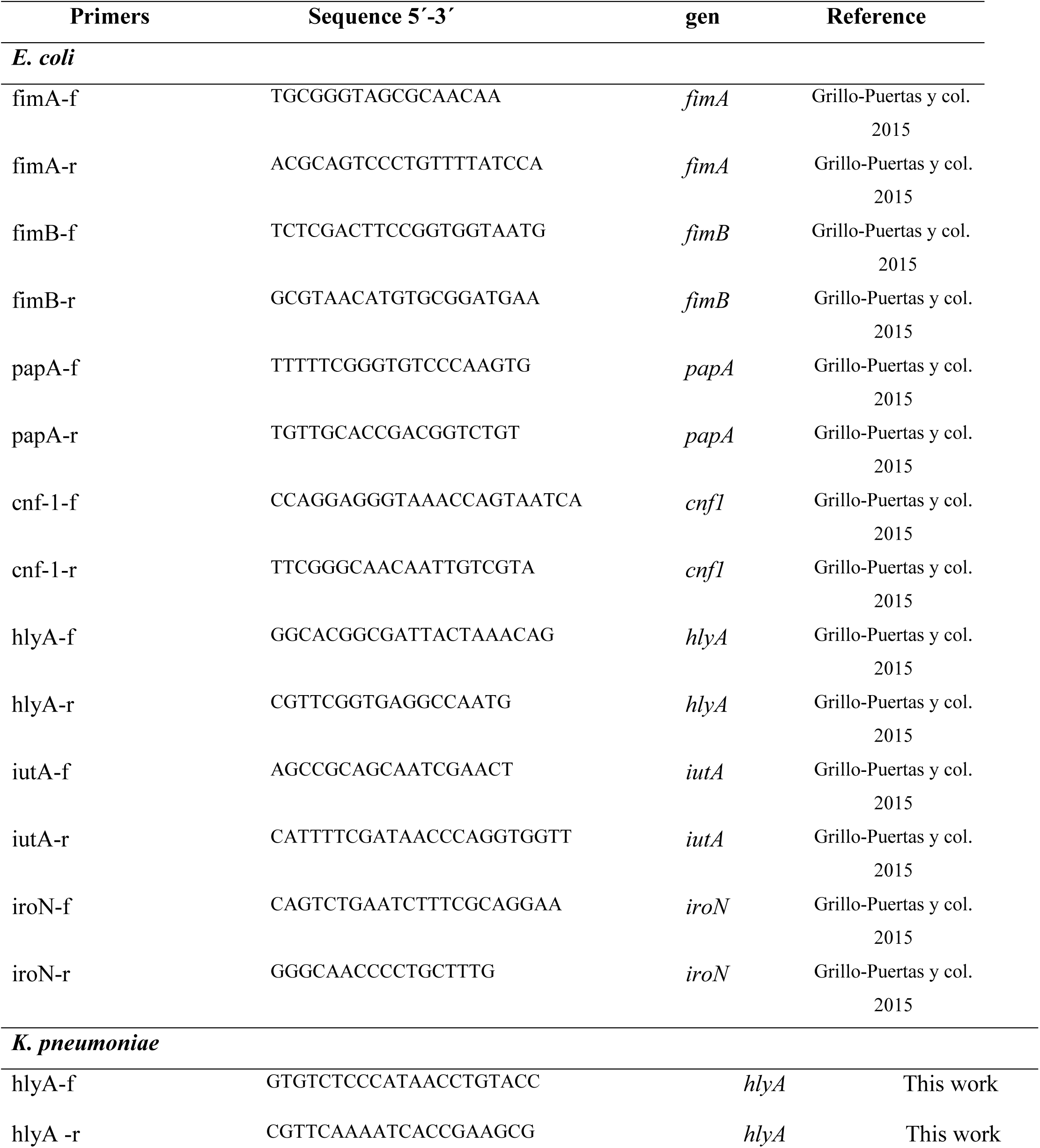
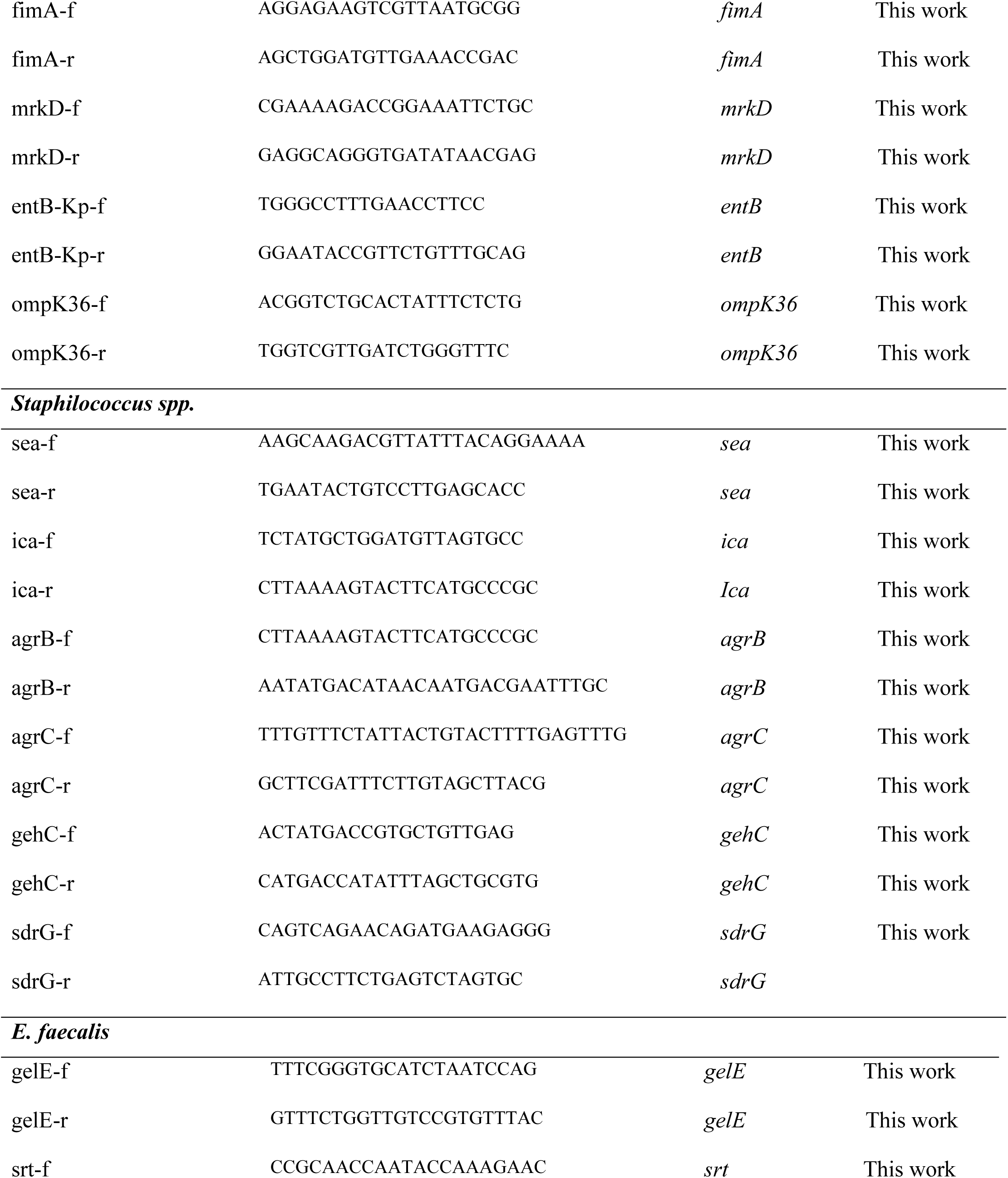
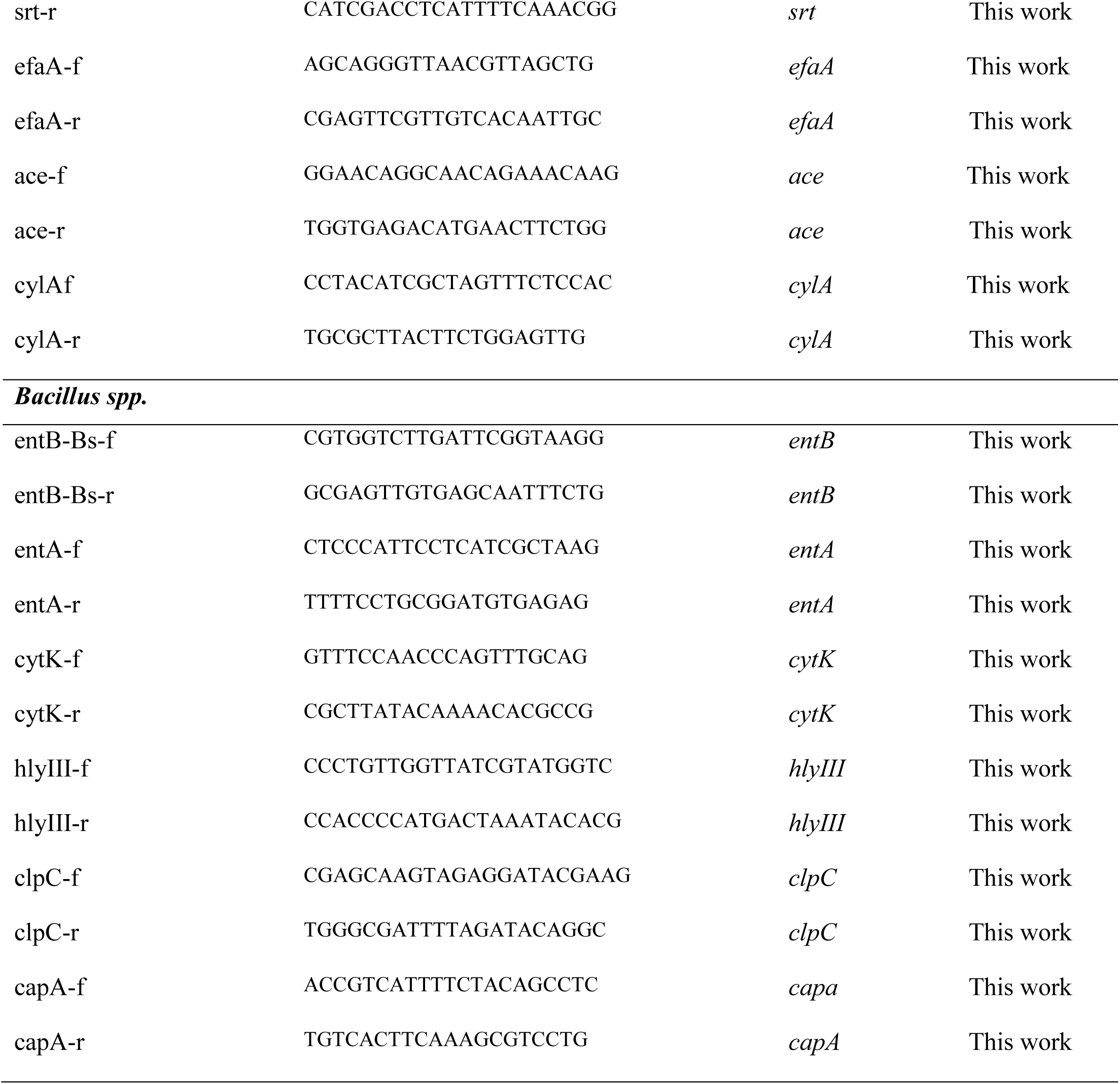
Primers used in this study.

**Figure S1:**
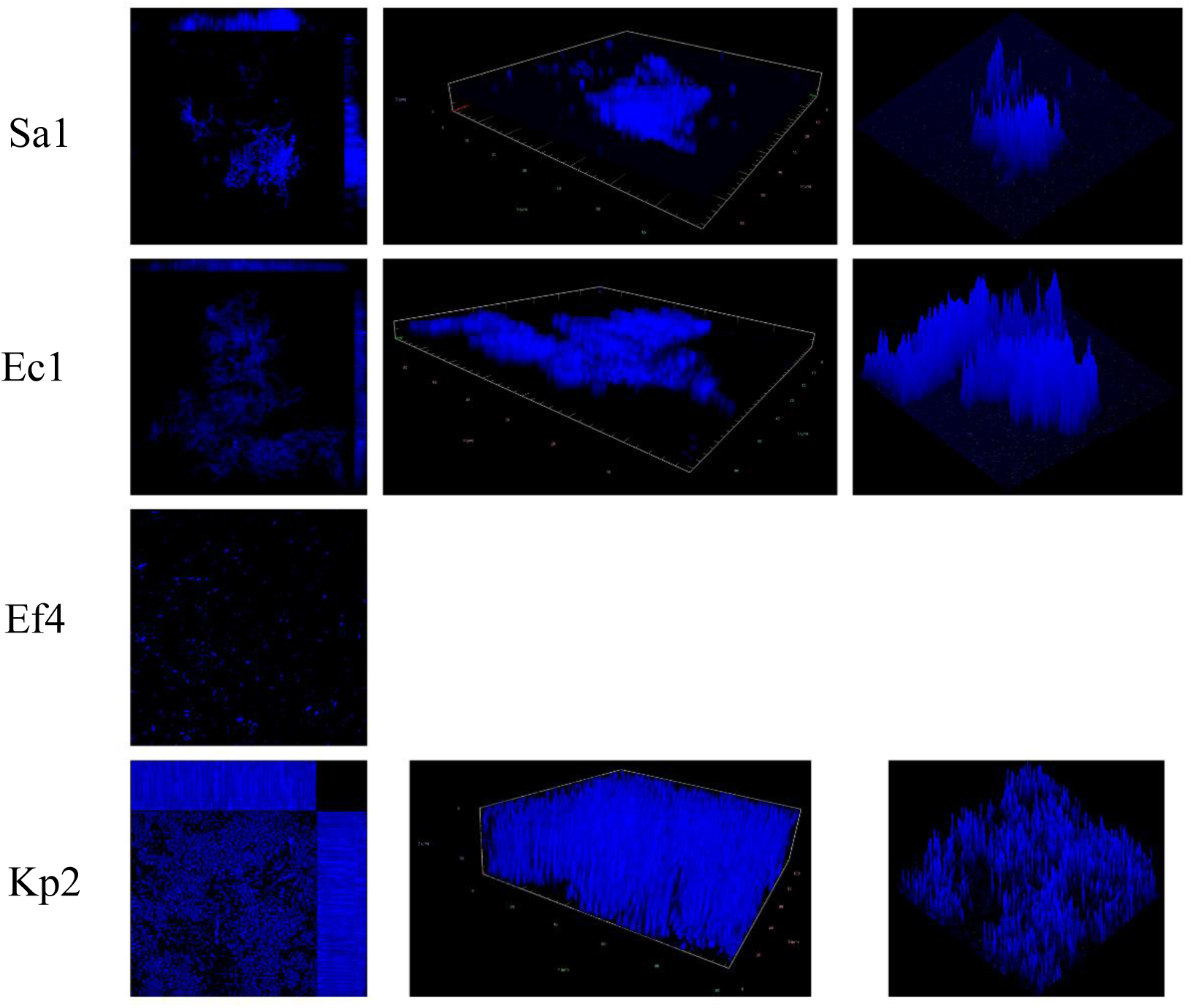
Laser scanning confocal microscopy analysis. The indicated isolates were grown in M63 medium for 72 h on specialized glass slides provided by the microscopy service. Subsequently, they were washed, and adhered cells were stained with DAPI (20 µM) for 15 minutes and fixed with paraformaldehyde. Biofilm formation was analyzed by CLSM (Zeiss LSM800), with a 63x immersion objective and the images represent three-dimensional reconstructions of the biofilm structures (ImageJ software). The images were representative of two independent experiments performed in duplicate.

**Figure S2:**
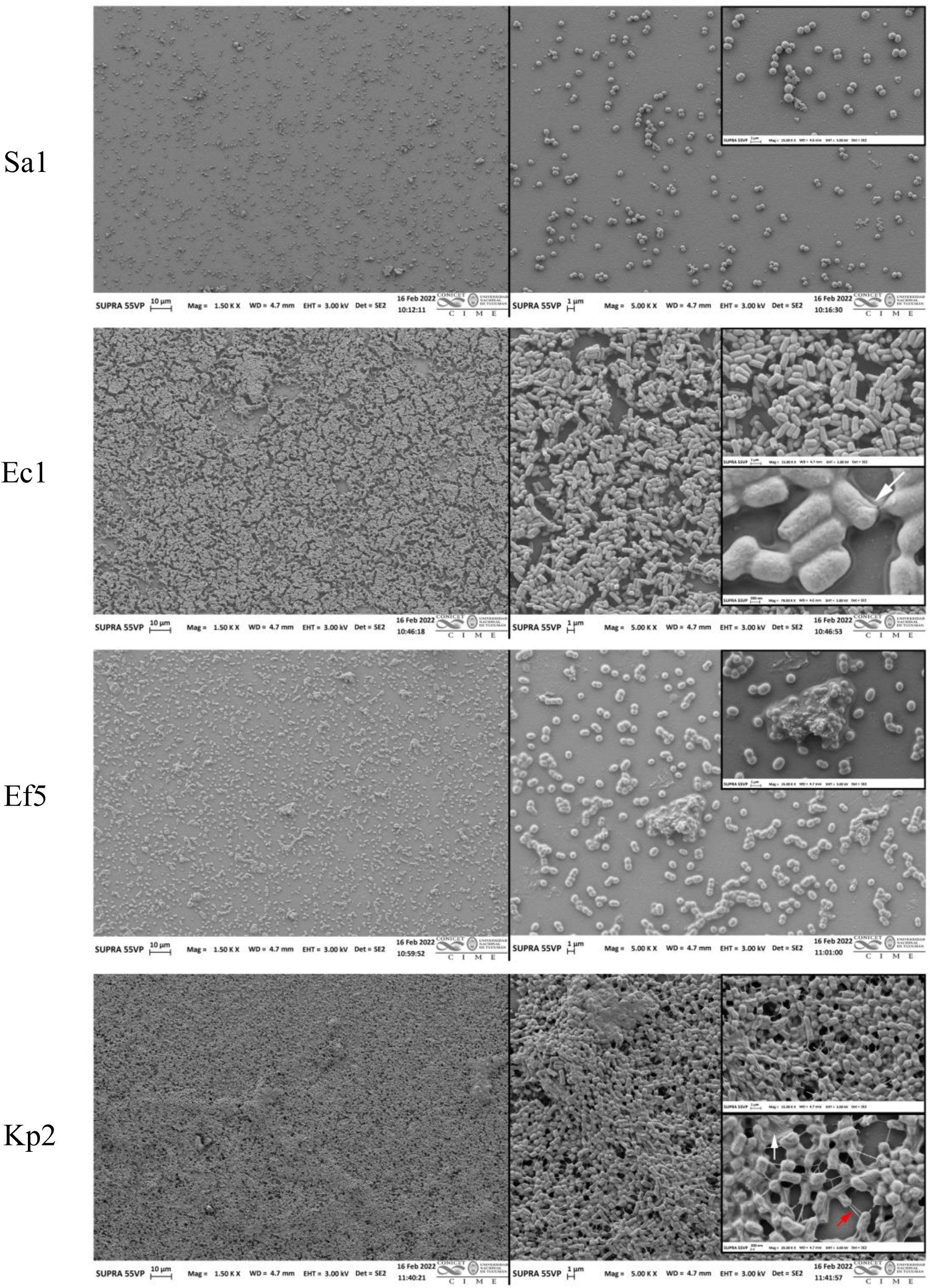
Scanning electron microscopy (SEM) analysis. The indicated isolates were grown in M63 medium for 72 h on specialized glass slides provided by the microscopy service. The slides were then washed, and adhered cells were fixed with glutaraldehyde. Biofilm formation was analyzed by SEM (Carl Zeiss SUPRA-55), with a resolution of 1.0 nm at 15 kV and 1.7 nm at 1 kV in HV (high-vacuum) mode and 2 nm at 30 kV in variable pressure (PV) mode. Magnification: Micrographs — left panels, 1500X; right panels, 5000X. Insert in EC1, 15000X; inserts in Ec1, 15000X and 70000X; insert in Ef5, 15000X; and inserts in Kp2, 15000X and 25000X. Images are representative of two independent experiments performed in duplicate.

**Figure S3.**
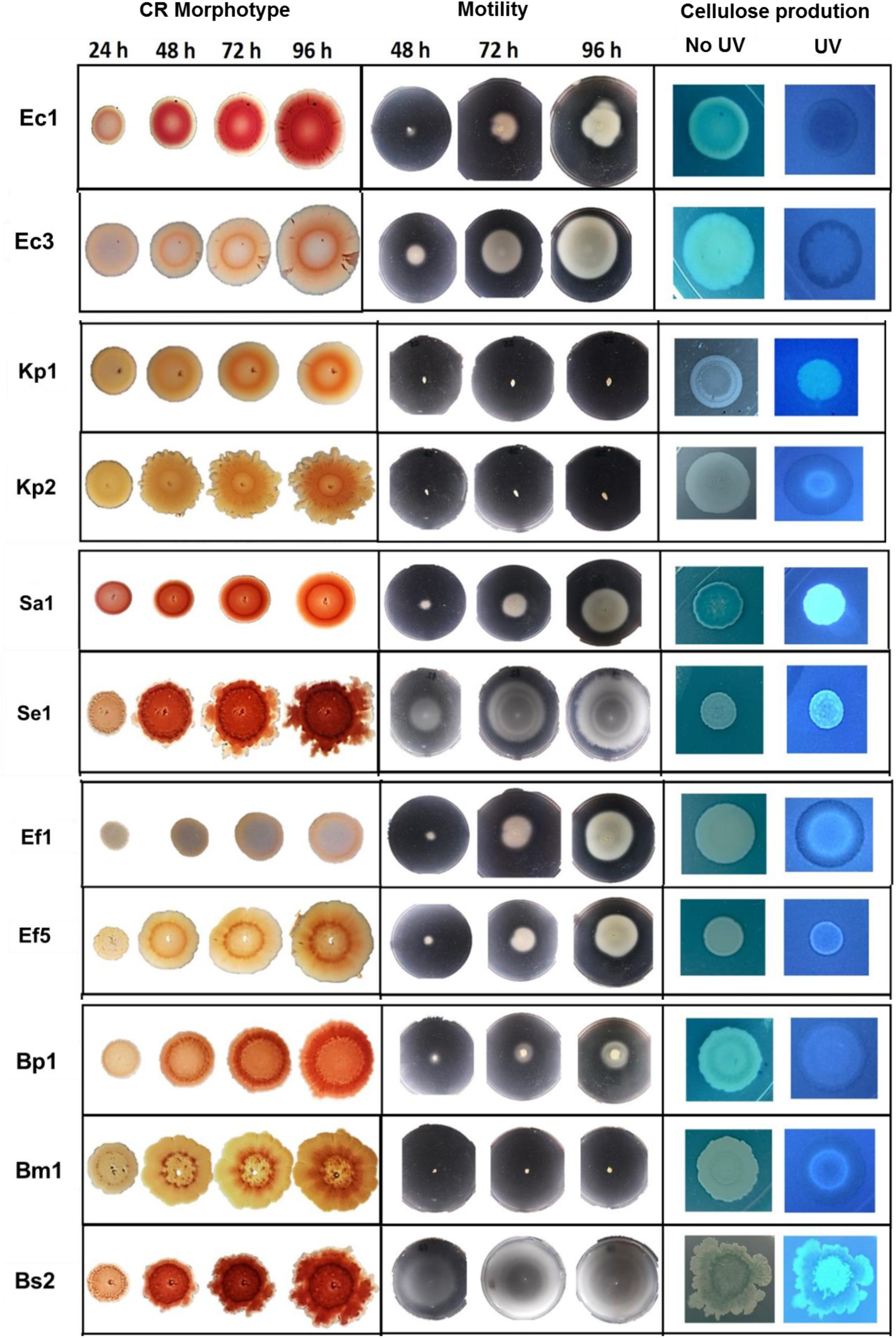
Bacterial virulence-associated phenotypes: *curli* or amyloid fibers, cellulose, and motility. Cultures of all isolates grown in LB or BHI medium for 24 h were washed and diluted to an A_600nm_ of 0.1. For morphology assays, suspensions were inoculated onto LB-agar plates (low Na^+^) supplemented with Congo Red (40 μg/ml) and Brilliant Blue (20 μg/ml). Motility analysis was carried out onto semi-solid M63-agar plates (0.3%) using a sterile toothpick for inoculation. Cellulose production was determined in LB-agar plates supplemented with Calcofluor White (50 μg/ml). Plates were incubated at 30°C during the indicated times for each assay and 48 h. Results are representative of four independent experiments.

**Figure S4.**
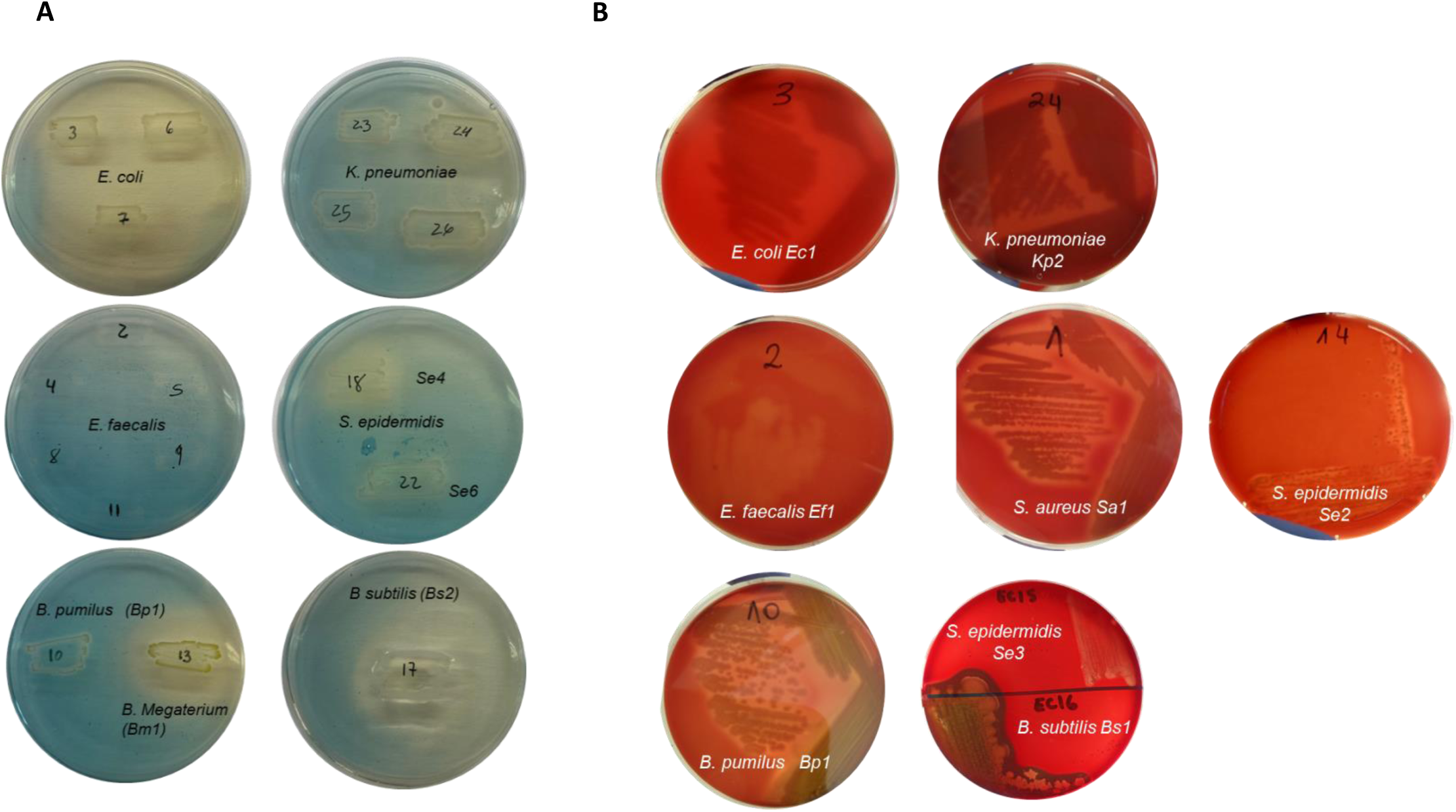
Siderophores production and hemolytic activity of the isolates. Siderophores production was evaluated by seeded isolates supentions on M9 minimal medium supplemented with 0.2% glucose plates and incubated at 37°C for 48 h. Then, an overlaid of semisolid CAS medium (see material and methods section) was poured onto the grown plate. Those isolates that are capable of producing siderophores presented a color change of the CAS medium from blue to yellowish around the colony (A). Hemolytic activity was evaluated on blood agar plates. Aliquots of bacterial suspensions were plated onto blood agar medium and incubated at 30°C for 24 hs. Hemolysis zones were interpreted as follows: -α hemolysis: partial lysis (turbid and greenish halo), -β hemolysis: complete lysis (clear, defined halo), -γ hemolysis: no hemolysis (B). Results are representative of three independent experiments.

**Table S2.**

| UP isolate |  | Antibiotic |  |  |  |  |  |  |  |  |  |  |  |  |  |  |  |  |
| --- | --- | --- | --- | --- | --- | --- | --- | --- | --- | --- | --- | --- | --- | --- | --- | --- | --- | --- |
|  |  | AMS | SXT | CIP | FT | CXM | CAZ | IMI | GM | AMP | NA | NOR | AKN | ERY | CLI | VAN | OXA | FOX |
| *E. coli* | Ec1 | R | R | R | S | S | S | S | S | R | R | R | S | - | - | - | - | - |
|  | Ec2 | R | R | R | S | S | S | S | S | R | R | R | S | - | - | - | - | - |
|  | Ec3 | R | R | R | S | S | S | S | R | R | R | R | S | - | - | - | - | - |
| *K. pneumoniae* | Kp1 | R | R | R | R | R | R | S | S | R | R | R | S | - | - | - | - | - |
|  | Kp2 | R | R | R | R | R | R | S | I | R | R | R | S | - | - | - | - | - |
|  | Kp3 | R | R | R | I | R | R | S | I | R | R | R | S | - | - | - | - | - |
|  | Kp4 | R | R | R | I | R | I | S | I | R | R | R | S | - | - | - | - | - |
| *S. aureus* | Sa1 | S | - | S | S | S | S | S | S | R | R | S | S | S | S | S | S | S |
| *S. epidermidis* | Se1 | S | - | S | S | S | S | S | S | R | R | S | S | R | S | S | S | S |
|  | Se2 | S | - | S | S | S | S | S | R | R | R | S | S | R | S | S | S | S |
|  | Se3 | S | - | S | S | S | S | S | R | R | R | S | S | R | R | R | S | S |
|  | Se4 | S | - | R | S | S | S | S | S | I | R | R | S | S | S | S | S | S |
|  | Se5 | S | - | S | S | S | S | I | S | I | R | S | S | S | S | S | S | S |
|  | Se6 | S | - | S | S | S | S | S | S | R | R | S | S | R | S | S | S | S |
| *E. faecalis* | Ef1 | I | - | R | S | R | R | S | R | S | R | R | S | R | R | S | R | R |
|  | Ef2 | S | - | R | S | R | R | S | R | S | R | R | R | R | R | S | R | R |
|  | Ef3 | S | - | R | S | R | R | S | R | S | R | R | I | R | R | S | R | R |
|  | Ef4 | S | - | R | S | R | R | R | R | S | R | R | S | R | R | S | R | R |
|  | Ef5 | S | - | R | S | R | R | R | R | S | R | R | R | R | R | S | R | R |
|  | Ef6 | S | - | R | S | R | R | S | R | S | R | R | I | R | R | S | R | R |
| *B. pumilus* | Bp1 | S | - | S | S | S | R | S | S | S | S | S | S | R | R | S | R | R |
| *B. megaterium* | Bm1 | S | - | S | S | S | S | S | S | R | S | S | I | S | R | S | S | S |
|  | Bm2 | S | - | S | S | S | S | S | S | S | S | S | S | S | S | S | S | S |
| *B. subtilis* | Bs1 | S | - | S | S | S | S | S | S | S | S | S | S | S | S | S | S | S |
|  | Bs2 | S | - | S | S | S | S | S | S | S | S | S | S | S | S | S | S | S |
|  | Bs3 | S | - | S | S | S | S | S | S | R | S | S | S | S | S | S | S | S |
|  | Bs4 | S | - | S | S | I | S | S | S | I | S | S | S | S | S | S | S | S |

**Figure S5.**
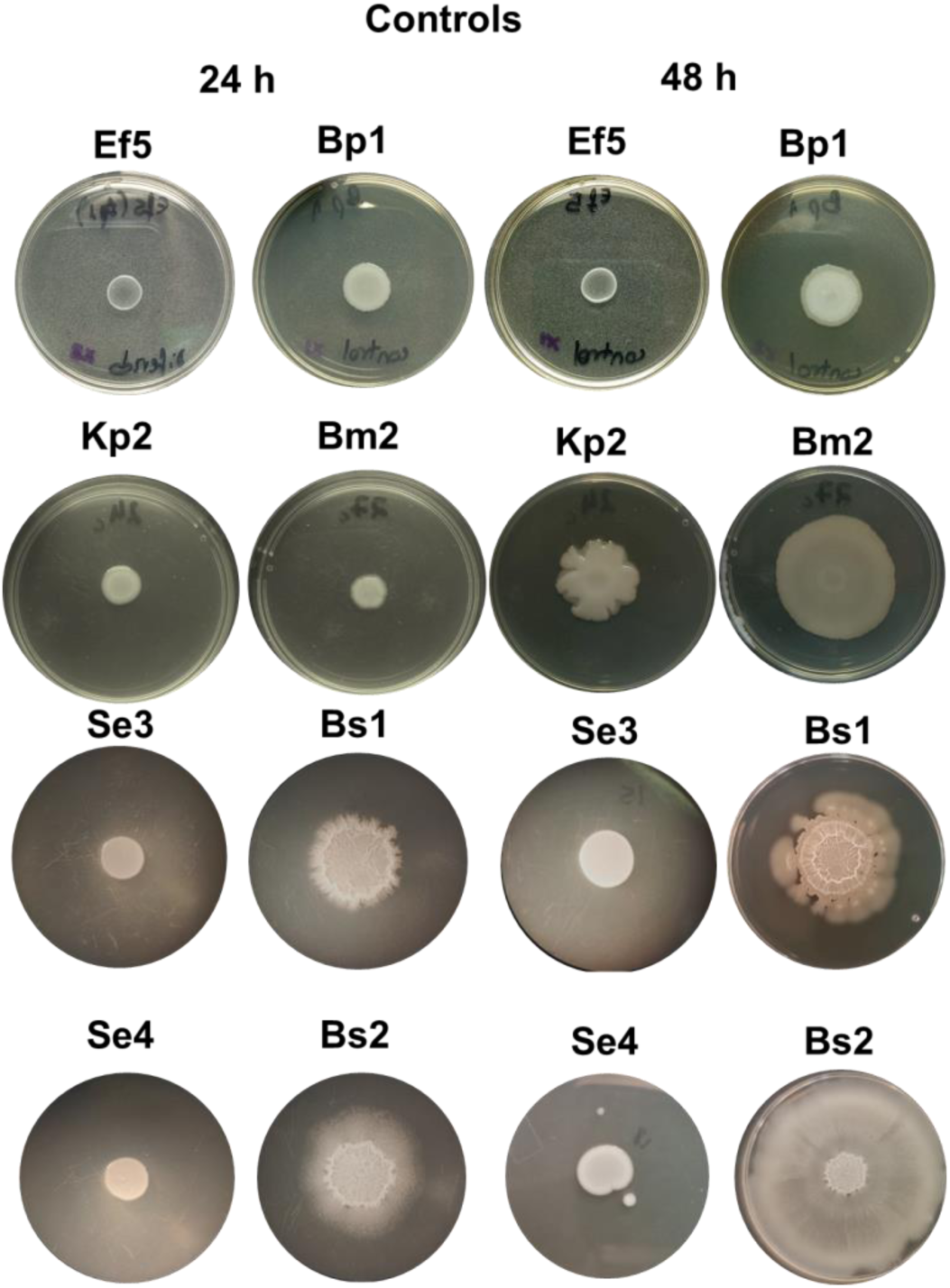
Single colonies incubation. 10 μl aliquots of the indicated isolates were spotted in the center of a BHI-agar plate and were grown up to the indicated times. Results represent four independent experiments.

